# Surprise gates two distinct mechanisms to support memorability in music

**DOI:** 10.64898/2026.07.21.739807

**Authors:** Mathieu Pham Van Cang, Paul Robert, Manuel Mercier, Agnès Trébuchon, Fabrice Bartolomei, Benjamin Morillon, Keith Doelling, Luc H Arnal

## Abstract

Music is a uniquely memorable human creation that, when skillfully composed, can persist in individual memory (as in *e.g.,* earworms) and cultural transmission (*e.g.* global hits or anthems). While both acoustic and statistical properties are known to influence a song’s memorability, the neural mechanisms that facilitate the engramming of certain musical sequences remain unclear. Current theories suggest that memory systems function as predictive internal models, enhancing learning when expectations are violated. Yet expected stimuli, by aligning with and reinforcing prior knowledge, also enhance memorability. How these two opposing processes arise from the brain’s sensitivity to statistical regularities, especially in naturalistic sequences, is not well understood.

Here, we leveraged music’s intrinsic balance between expectancy and surprise, and examined the neural correlates of minutes-scale memorability using intracranial EEG recordings from nine patients with epilepsy performing a musical memory task. Quantifying the statistical surprise of each musical excerpt with PolyRNN, a polyphonic model of musical expectations, we uncovered a U-shaped relationship between musical surprise and memory performance: both highly expected and highly surprising melodies led to greater memorability. While neural pattern similarity between song repetitions was enhanced for low-surprise stimuli, high-surprise stimuli enhanced neural pattern separability in medial temporal regions, each mediating memorability in distinguishable ways. These findings reveal complementary neural mechanisms through which statistical structure shapes musical sequence memory, clarifying how the brain encodes complex, ecologically valid stimuli.

## Introduction

Not all sensory experiences are equally memorable, and music stands out as a striking example of a stimulus that imprints itself in both individual memory and cultural embedding. Songs and melodies often linger involuntarily in our minds, as earworms (*1*), or stand as powerful cultural symbols, from anthems to global hits (*2*, *3*). Memorability is not merely a byproduct of musical experience but a key driver of cultural selection: the most memorable songs are more likely to endure and spread. To achieve lasting impact, composers balance *(i)* expectancy, aligning with listeners’ prior experience and knowledge, and *(ii)* surprise, introducing unexpected elements that capture attention. These two components, as well as their balance, influence how music evokes emotion (*4*, *5*). While various acoustic and statistical features may contribute to a song’s memorability, the neural mechanisms that support the encoding of musical sequences remain poorly understood.

Current theories propose that neural memory systems function as internal models of the world (*6*), contributing to shaping our predictions about incoming sensory information. When these predictions differ from actual inputs, the mismatch provides crucial learning signals that enhance the encoding of newly encountered patterns. For instance, unexpectedly encountering a complex chord progression in a piece of music that otherwise shows a simple structure might sharpen memory for that moment. Conversely, stimuli that align with prior expectations may also be easier to remember, as they fit seamlessly into existing cognitive schemas (*e.g.*, familiar melodies or frequently rehearsed tunes recalled effortlessly). These seemingly contradictory effects reflect the brain’s sophisticated use of statistical regularities to optimize the storage of complex sequences in memory. Indeed, while surprising events often capture attention and can be better remembered (*7*), the precise interplay between surprise, expectancy, and memory remains elusive.

Music offers an ecologically rich framework to investigate these dynamics (*8*). Listening to music engages predictive processing: the brain draws on learned statistical patterns to anticipate upcoming sounds (*4*, *5*). Violations of these predictions, such as a sudden key change or a rhythmic disruption, generate prediction errors that update the brain’s internal model (*9–11*) and enhance memory encoding (*12*). This aligns with findings showing that novel or surprising sequences are often better remembered, highlighting the brain’s sensitivity to surprise (*7*, *13*). At the same time, expected patterns also show memory advantages, likely because they resonate with established representations and are reinforced through repeated exposure (*14*). This creates a paradox: both surprise and predictability appear to support memory.

Empirical studies using artificial sequences have documented memory benefits from both surprise and predictability (*15–17*). However, contradictory findings (*18*, *19*) and the lack of a mechanistic framework leave open questions about how these effects co-exist. Particularly, it is unlikely that a single neural mechanism can fully account for two opposite effects: the enhanced encoding of both high- and low-surprise events.

Here, we hypothesized that surprise modulates two distinct neural mechanisms to facilitate memory performance: *(i) pattern similarity* for low-surprise (expected) events, reflected by a stronger overlap in neural responses to distinct presentations of the same stimulus (*20–24*), and *(ii) pattern separability* for high-surprise (unexpected) events, reflected by a greater neural differentiation between stimuli (*24–28*). By disentangling these complementary processes, we aim to clarify how statistical regularities shape memory for complex sequences like music.

Such interactions between predictions and memory are likely to manifest in the medial temporal lobe. Prior research has demonstrated that the hippocampus and adjacent regions are sensitive to auditory stimuli (*29–31*) and encode statistical regularities in sound sequences (*32–34*). Moreover, these areas are implicated in the detection of both novel (*35–40*) and familiar stimuli (*41–43*). How these processes interact and whether they are encoded in overlapping or distinct circuits remain open questions.

To address these questions, we developed a naturalistic musical memory experiment using an old/new recognition task with 9 epilepsy patients undergoing presurgical evaluation via stereotactic EEG recordings. We analyzed both behavioral performance and its potential neural correlates using a statistical model-based approach, focusing on temporal lobe responses.

The stimuli consisted of excerpts from classical polyphonic piano music, chosen for their rich structural complexity. For each excerpt, we calculated trial-by-trial surprise values using *PolyRNN*, a recurrent neural network trained to model tonal and temporal polyphonic musical expectations (*44*). This computational approach enabled us to probe how surprise modulates neural pattern similarity and separability, and how these neural signatures relate to memory outcomes. Specifically, following the memory literature (*45*), we tested the prediction that low-surprise clips would elicit greater neural response similarity across repetitions, while high-surprise clips would evoke more distinct and separable neural patterns within the medial temporal lobe.

## Results

### Experimental setup and hypothesis

Patients listened to 8-second excerpts of classical polyphonic piano performances, each presented twice during the experiment (**Fig. 1A**). The task followed an old/new recognition paradigm: participants indicated, via button press, whether each clip was “old” (previously heard) or “new”. The presentation order was pseudo-randomized to ensure a balanced distribution of new and old trials (see **Methods**).

**Figure 1:**
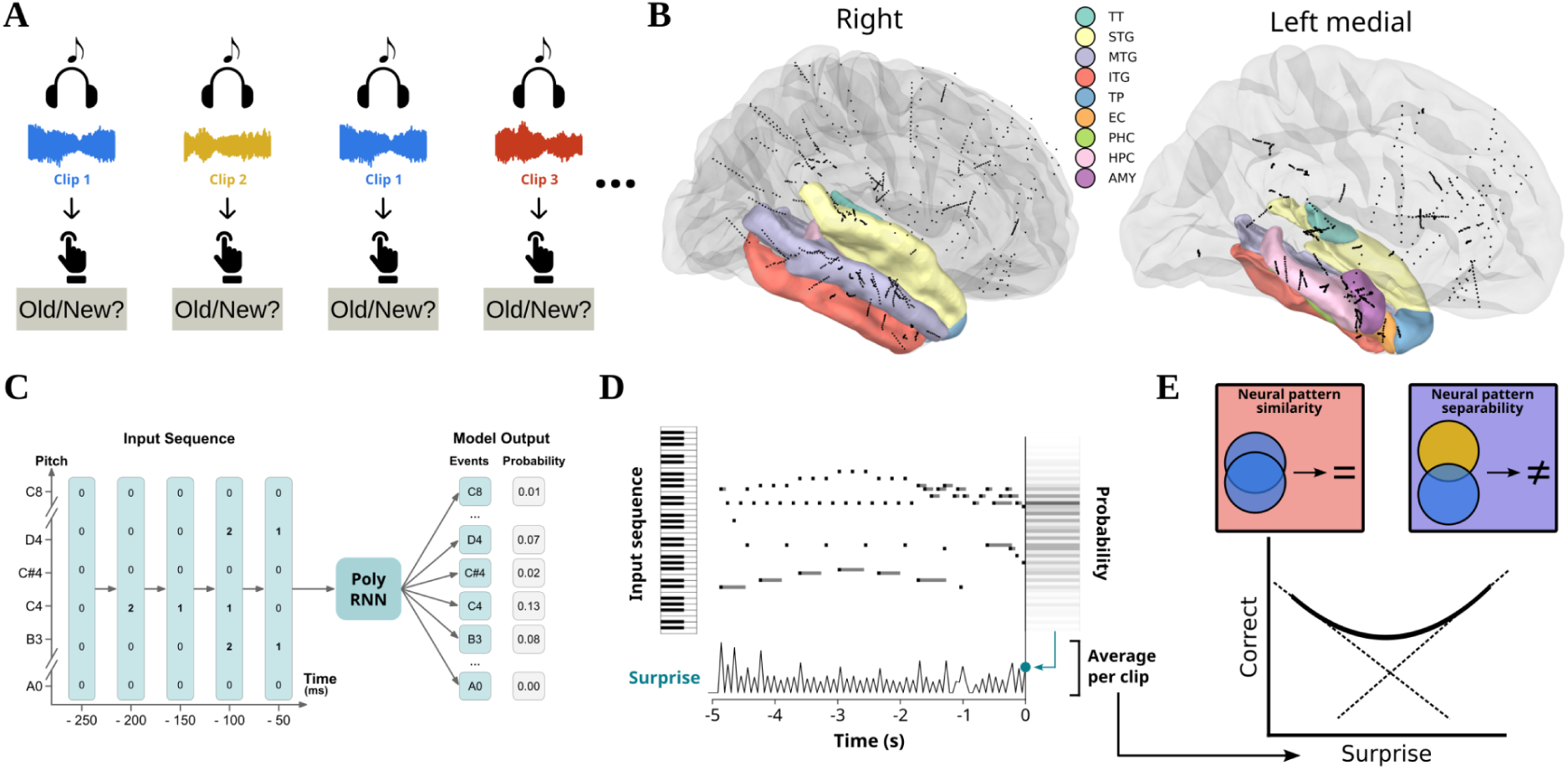
Experimental design and hypotheses. **A.** Old/new recognition paradigm. **B.** sEEG channel coverage across 9 patients and parcellation of the temporal lobe regions (n = 291 channels) included in the analyses (9 regions per hemisphere, based on MNE’s aparc+aseg segmentation (*46*, *47*). **C.** Architecture of the *PolyRNN* deep learning model for musical prediction. Once trained, the model outputs note probabilities at each time step based on preceding musical context. **D.** Note probabilities are then used to compute a time series of surprise values. For each clip, the average surprise over time is calculated and used in all current analyses. **E.** Hypothesized nonlinear (U-shape) influence of musical contextual surprise on memory performance via two distinct neural processes: low-surprise clips elicit responses that are more stable across repetitions (higher pattern similarity), enhancing recognition, whereas high-surprise clips generate stronger prediction error signals that are easily discriminable across clips (higher pattern separability), reducing confusion with previously heard clips, which also enhances recognition. TT: transverse temporal, STG: superior temporal gyrus, MTG: middle temporal gyrus, ITG: inferior temporal gyrus, TP: temporal pole, EC: entorhinal cortex, PHC: parahippocampal cortex, HPC: hippocampus, AMY: amygdala.

The 9 patients were implanted with a total of 134 sEEG electrode shafts (median: 15, range: 12-17), resulting in 1479 channels. **Figure 1B** shows full channel coverage and highlights the 9 regions of interest, all located within the temporal lobes (n = 291 channels, both hemispheres are included for each region, see **Methods** and **Table S1** for parcellation methods and detailed channel coverage). Signal analyses focused exclusively on channels located in these regions.

For each excerpt, we quantified musical statistical surprise using *PolyRNN*, a deep learning model that estimates tonal and temporal expectations in polyphonic music in a temporally-resolved manner (*44*). This model provides a *surprise* value for each note or chord, reflecting the unexpectedness of each musical event. For each clip, we computed an average surprise score across all notes to derive a trial-level measure of statistical surprise (**Fig. 1C-D**; see **Methods**).

We hypothesized that surprise would exert a nonlinear influence on memory performance, mediated by two complementary neural mechanisms: *pattern similarity*, which supports recognition by increasing overlap in neural responses across repetitions of the same clips, and *pattern separability*, which reduces interference by enhancing neural distinctiveness between clips (**Fig. 1E**; see **Methods**). This framework predicts that recognition of low-surprise clips would benefit from enhanced similarity, while high-surprise clips would be encoded more distinctly through enhanced separability, both neural mechanisms facilitating memorability.

### Memory performance in novelty detection and recognition trials

We first examined the behavioral performance on the old/new recognition task, separating trials by their presentation rank to assess both novelty detection (1^st^ presentations; correct response: “new”) and recognition (2^nd^ presentations; correct response: “old”). Participants performed the task reliably, showing above chance accuracy in both novelty trials (one-sample one-tailed *t*-test: *t*(8) = 6.85, *p* < 0.001; **Fig. 2B**) and recognition trials (one-sample one-tailed *t*-test: *t*(8) = 2.68, *p* = 0.014; **Fig. 2C**), as well as a difference in response between novelty and recognition (paired two-tailed *t*-test: *t*(8) = 6.33, *p* < 0.001).

**Figure 2:**
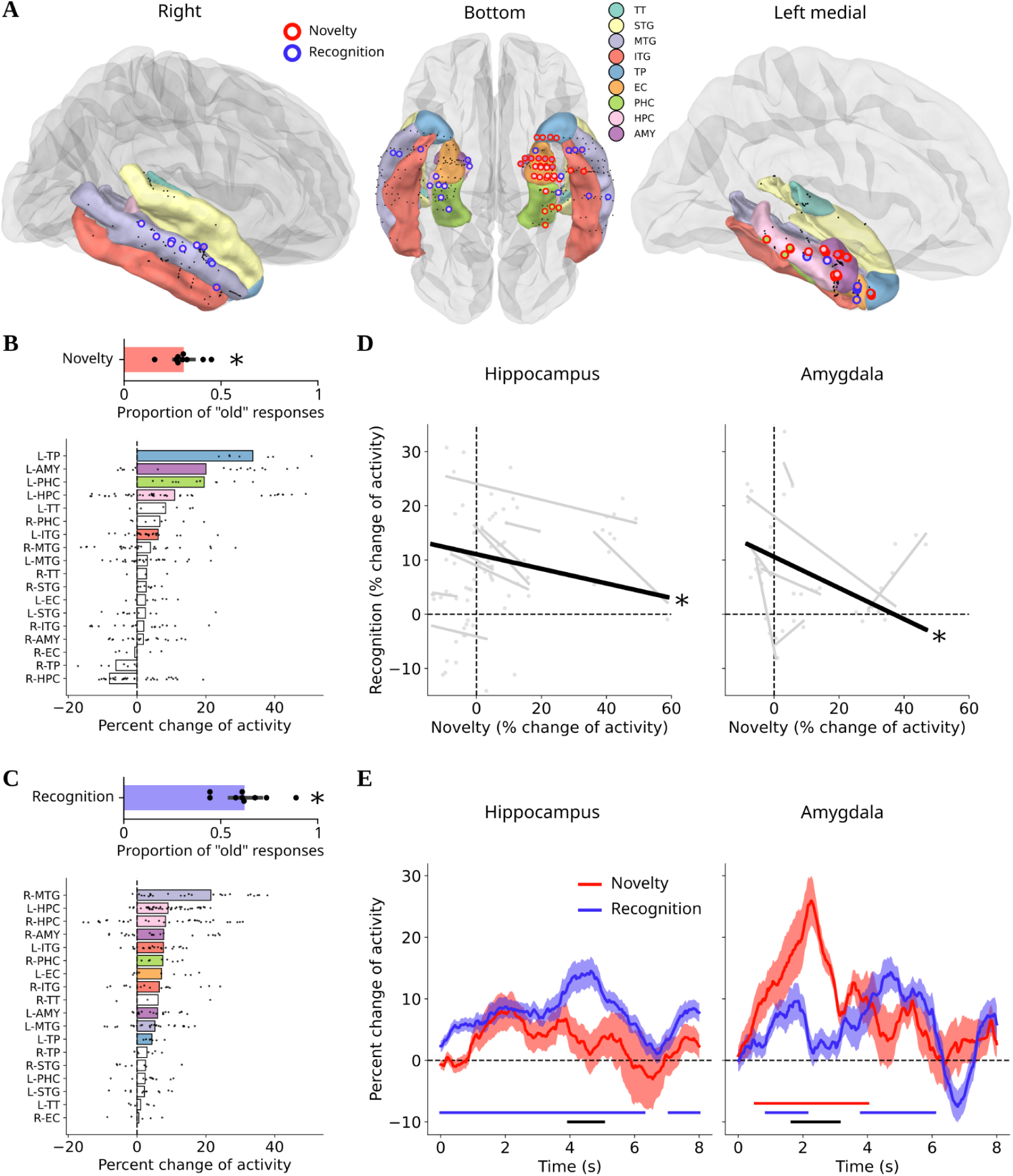
Memory performance is associated with increased neural activity in the temporal lobe. **A.** Channels showing a significant activity contrast (correct minus incorrect trials, permutation-based *t*-tests, *p* < 0.05) in novelty detection (red) and recognition (blue) trials. No spatial overlap between effects was observed. Inner colors indicate the anatomical location of the channels. **B.** Top: proportion of “old” responses in novelty detection trials. Subjects (n = 9) are represented by black dots. Behavior was significantly better than chance level (*t*-test, * *p* < 0.05). Bottom: bar plot of activity contrasts by region in novelty detection trials. Black dots represent individual channels. Filled colored bars indicate regions with a significant contrast (*t*-tests, FDR-corrected across regions, *p* < 0.05). **C.** Same as B in recognition trials. **D.** Recognition activity contrast plotted as a function of novelty activity contrast in the hippocampus (left, n = 77 channels) and amygdala (right, n = 39 channels). Gray dots/lines represent individual channels/subjects; the black line shows the average relationship (fixed effect from a GLMM, * *p* < 0.05). **E.** Activity contrasts over time in novelty detection and recognition, averaged across channels in the hippocampus (left, n = 77) and amygdala (right, n = 39). Shaded areas represent the SEM across channels. Lines at the bottom indicate significant temporal clusters of activity (red and blue), or differences between novelty and recognition (black), obtained from permutation-based temporal cluster *t*-tests (*p* < 0.05). TT: transverse temporal, STG: superior temporal gyrus, MTG: middle temporal gyrus, ITG: inferior temporal gyrus, TP: temporal pole, EC: entorhinal cortex, PHC: parahippocampal cortex, HPC: hippocampus, AMY: amygdala, L: left hemisphere, R: right hemisphere.

### Memory performance is associated with increased neural activity in the medial temporal lobe

Next, we hypothesized that neural activity in the temporal lobe, specifically within the medial temporal lobe, would be associated with correct memory performance. We therefore focused our analysis on channels located within the temporal lobe (**Fig. 1B**). We used the trial-level activity of the intracranial recordings (see **Methods**), sorted as a function of behavioral data to contrast correct versus incorrect trials. This analysis was conducted separately for novelty detection and recognition trials. This revealed channels that showed a significant activity contrast in either novelty detection or recognition (*p* < 0.05; two-sample permutation-based *t*-tests across trials; **Fig. 2A**), with no channel showing the effect in both processes. This analysis also showed a left-lateralized pattern of activation during novelty detection (30 significant channels out of 160 in the left hemisphere, none out of 131 in the right hemisphere), while recognition did not show such lateralization (8 significant channels in the left hemisphere, 11 in the right hemisphere).

To further identify which temporal lobe regions exhibited these effects, we ran statistical tests across channels within each anatomical region. We found a significant increase in neural activity during correct trials in several regions (two-tailed paired *t*-tests with FDR correction across regions). For novelty detection (**Fig. 2B**), significant effects were observed in 5/18 regions: the left amygdala (*t*(17) = 4.49, *p* = 0.002), left temporal pole (*t*(6) = 9.31, *p* = 0.001), left parahippocampal gyrus (*t*(11) = 5.74, *p* = 0.001), left inferior temporal gyrus (*t*(25) = 2.67, *p* = 0.047), and left hippocampus (*t*(42) = 3.62, *p* = 0.004). For recognition (**Fig. 2C**), significant effects were found in 11/18 regions: the left and right amygdala (*t*(17) = 4.91, *p* < 0.001; *t*(20) = 2.88, *p* = 0.019), left temporal pole (*t*(6) = 5.60, *p* = 0.005), left entorhinal cortex (*t*(8) = 2.77, *p* = 0.040), right parahippocampal gyrus (*t*(7) = 4.91, *p* = 0.005), left and right middle temporal gyri (*t*(26) = 3.19, *p* = 0.008; *t*(25) = 4.06, *p* = 0.002), left and right inferior temporal gyri (*t*(25) = 4.71, *p* < 0.001; *t*(17) = 2.55, *p* = 0.037), and left and right hippocampi (*t*(42) = 9.41, *p* < 0.001; *t*(33) = 3.39, *p* = 0.005). These results confirm a left-lateralized network for detecting novel musical pieces and highlight the involvement of a broader, bilateral network in the recognition of previously heard melodies. Overall, increased neural activation in higher-level auditory regions and the medial temporal lobe was associated with better memory performance.

### Novelty detection and recognition are supported by distinct neural populations

We then tested whether the neural populations supporting novelty detection and recognition overlapped or were entirely distinct. To test this, we examined the relationship between novelty detection and recognition trials in terms of the performance-related increase of activity (activity contrast between correct and incorrect trials) across channels. Our rationale was that if these two processes depend on distinct neural populations within regions, then we should find a negative relationship in the activity contrast. For example, if a channel is close to a subpopulation specifically involved in novelty detection, it will be farther from subpopulations involved in recognition. We focused the analysis on the hippocampus and amygdala (see **Fig. S1** for the same analysis on other regions of the temporal lobe), as these regions had the largest channel coverage and have been shown to contribute to old/new memory paradigms (*48*, *49*). We observed a negative relationship between the two types of trials in both the hippocampus (GLMM: β = −0.13 ± 0.04 SE, *z* = −3.23, *p* = 0.001) and the amygdala (β = −0.29 ± 0.11 SE, *z* = −2.62, *p* = 0.009; **Fig. 2D**). This confirms that, although these regions are involved in both novelty detection and recognition, the underlying neural populations supporting each process are distinct (**Fig. 2A**). We did not observe any average difference in neural activity between trials followed by a “new” response and trials followed by an “old” response, ruling out the possibility that our results were the consequence of a difference in activity independently of memory performance.

Next, we asked whether novelty detection and recognition are associated with similar or distinct neural dynamics over the course of the musical excerpts. We contrasted neural activity between correct and incorrect trials in a time-resolved manner, using a 1-second rolling window across the 8-second clips to examine the temporal evolution of the performance-related neural activity in both trial types. Again, we focused on the hippocampus and amygdala (**Fig. 2E**). We first tested activity contrast against 0 in each condition through permutation-based temporal cluster two-tailed *t*-tests across channels. In the hippocampus, we found two significant clusters of activity contrast during retrieval (*t*-threshold = 1.99, *p*_1_ < 0.001, *p*_2_ = 0.038). In the amygdala, we found one significant cluster during novelty (*t*-threshold = 2.02, *p* < 0.001) and two significant clusters during retrieval (*t*-threshold = 2.02, *p*_1_ = 0.026, *p*_2_ < 0.001). We then tested for differences between novelty and recognition dynamics, and found a late period in which performance-related activity was significantly stronger for recognition in the hippocampus (permutation-based temporal cluster two-tailed, paired-sample *t*-test; *t*-threshold = 1.99, *p* = 0.044). In contrast, we found an early period with stronger activity for novelty in the amygdala (*t*-threshold = 2.02, *p* = 0.005). In novelty trials, the performance-related activity in the amygdala rose rapidly and gradually returned to baseline, suggesting that novelty detection is a fast process occurring primarily within the first few seconds of a clip. In recognition, we observed a ramping of the activity over time in the hippocampus, consistent with a more gradual accumulation of evidence throughout the listening period (*50*). Other regions of the temporal lobe showed varying patterns (**Fig. S2**).

Taken together, these results suggest that novelty detection and recognition are complementary processes supported by overlapping regions but rely on distinct neural populations, building evidence that novel and previously encoded stimuli engage different neural dynamics.

Throughout the rest of this paper, we restrict our analyses to brain regions that showed a significant activity contrast in either novelty (**Fig. 2B**) or recognition (**Fig. 2C**) trials, assuming that these regions were involved in this memory task. We call them regions of interest (ROIs) in all following sections. This process resulted in twelve selected regions (7/12 in the left hemisphere): middle temporal gyrus, inferior temporal gyrus, left temporal pole, left entorhinal cortex, parahippocampal cortices, hippocampus, amygdala.

### Neural pattern similarity and separability jointly support memory in the temporal lobe

To further investigate the neural mechanisms involved in memory, we studied two potential mechanisms: neural *pattern similarity* and *pattern separability*, hypothesizing that both would influence memory performance in this task (**Fig. 1E**). We used the correlation between the neural responses to the two presentations of a given stimulus as a measure of pattern similarity (**Fig. 3A**). On the other hand, we used the distance between the neural response to a given stimulus and the responses to all other previously heard clips as a measure of pattern separability (**Fig. 3B**; see **Methods**).

**Figure 3:**
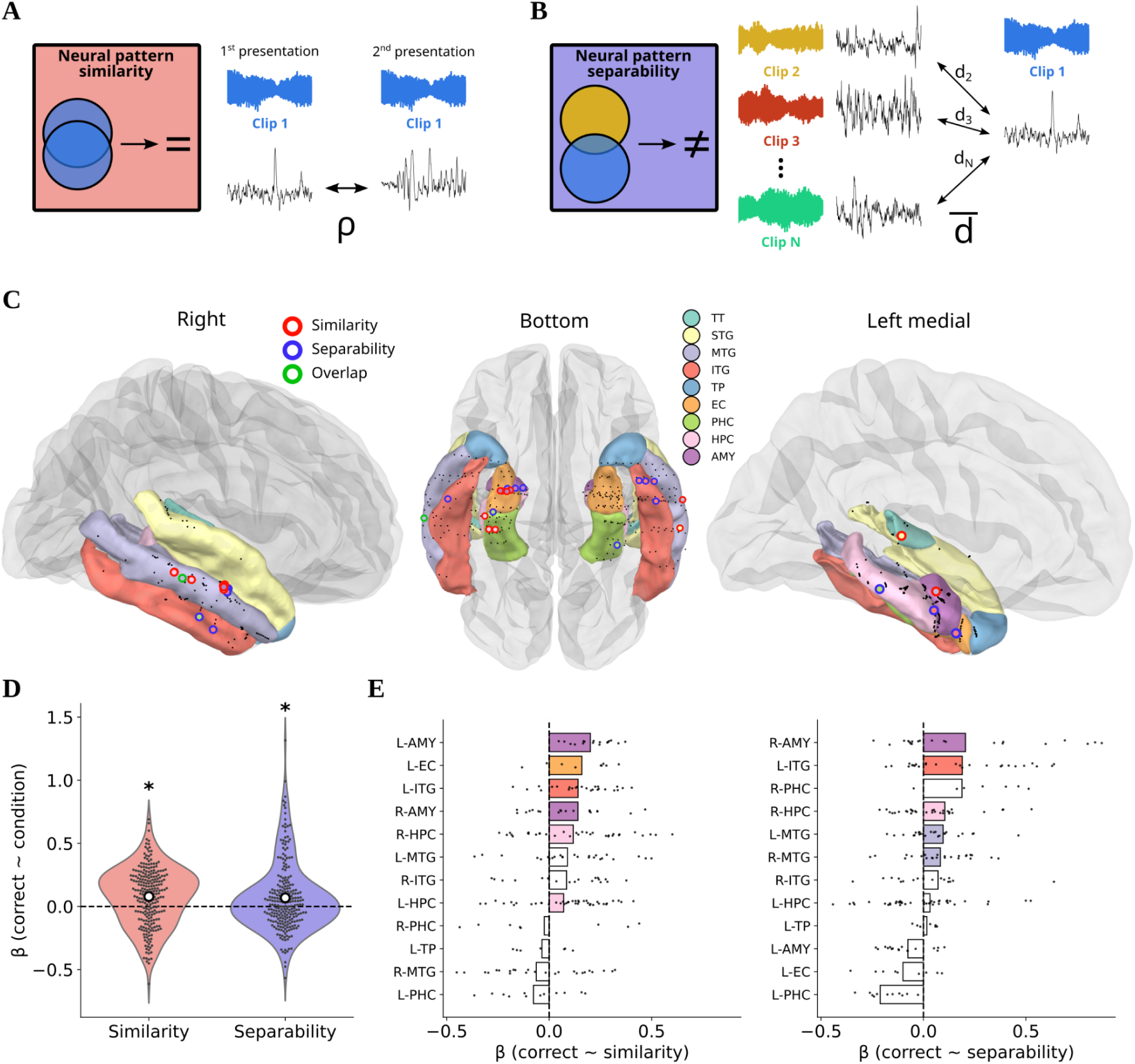
Distinct mechanisms of pattern similarity and separability jointly support memorization. **A.** Neural pattern similarity, measured as the Pearson’s correlation between the neural responses to the two presentations of a clip. **B.** Neural pattern separability, measured as the average distance between the neural response to a given clip and the responses to all other clips. **C.** Channels showing a significant relationship (logistic regression parameter, *p* < 0.05) between memory accuracy and similarity (red) or separability (blue). One channel showed both effects and is indicated in green. Inner colors indicate the anatomical location of the channels. **D.** Violin plots of logistic regression parameters in the ROIs (n = 249 channels) for the relationships *correct ∼ similarity* and *correct ∼ separability* (*t*-test, * *p* < 0.05). White dots indicate the mean. **E.** Bar plot of logistic regression parameters by region for the two relationships (left: similarity, right: separability). Black dots represent individual channel parameters. Filled colored bars indicate regions with a significant positive average (*t*-tests, FDR-corrected, *p* < 0.05). TT: transverse temporal, STG: superior temporal gyrus, MTG: middle temporal gyrus, ITG: inferior temporal gyrus, TP: temporal pole, EC: entorhinal cortex, PHC: parahippocampal cortex, HPC: hippocampus, AMY: amygdala, L: left hemisphere, R: right hemisphere.

We first assessed the influence of these neural pattern measures on memory using subjects’ responses in both novelty and recognition trials. We predicted trial-level memory performance from either similarity or separability values, for each channel of the ROIs (logistic regressions across trials; *p* < 0.05; **Fig. 3C**). We tested the robustness of such effects across channels in the ROIs (one-tailed one-sample *t*-test on the regression parameters), and confirmed a significant positive linear relationship between correct responses and both similarity (*t*(248) = 6.00, *p* < 0.001) and separability (*t*(248) = 4.37, *p* < 0.001; **Fig. 3D**). This indicates that both a stronger pattern similarity between two presentations of the same clip and a stronger pattern separability between a clip and all the previously heard excerpts positively impact memorability.

We next investigated the regional specificity of these effects (one-tailed one-sample *t*-tests with FDR correction for the number of regions). We found a significant effect of similarity on memory accuracy in 6/12 regions: the left and right amygdala (*t*(17) = 7.93, *p* < 0.001; *t*(20) = 2.61, *p* = 0.021), the left entorhinal cortex (*t*(8) = 3.03, *p* = 0.021), the left inferior temporal gyrus (*t*(25) = 4.12, *p* = 0.001), the left and right hippocampus (*t*(42) = 2.12, *p* = 0.04; *t*(33) = 2.74, *p* = 0.02; **Fig. 3E**, left). For separability, we found a significant effect in 5/12 regions: the right amygdala (*t*(20) = 2.60, *p* = 0.042), the left inferior temporal gyrus (*t*(25) = 3.73, *p* = 0.006), the right hippocampus (*t*(33) = 2.30, *p* = 0.042), the left and right middle temporal gyrus (*t*(26) = 2.41, *p* = 0.042; *t*(25) = 2.15, *p* = 0.049; **Fig. 3E**, right).

To ensure that the observed effect was not simply due to a genuine positive correlation between our two neural pattern measures, we directly correlated similarity and separability in channels that showed a significant relationship with memory performance in at least one of the two regressions. We found no significant correlation across epochs (GLMM: β = 0.009 ± 0.036 SE, *z* = 0.244, *p* = 0.807; see **Methods**), ruling out this possibility and confirming that similarity and separability, while distinct, both improve memory. Further visualizations of the joint effects of similarity and separability on memory performance are available in **Fig. S3**.

### Musical surprise exerts a U-shaped influence on memory

Next, we investigated the influence of statistical surprise on memory performance in the behavioral data. We fitted a Generalized Linear Mixed Model (GLMM) to predict trial-level memory accuracy (correct/incorrect). As we hypothesized that both low- and high-surprise clips would be associated with better memory performance (see **Fig. 1E**), we included squared surprise as a predictor in our model (along with control predictors, see **Methods** for modeling details). The effect of squared surprise on memory performance was significant (β = 0.11 ± 0.05 SE, *z* = 2.42, *p* = 0.015), revealing a U-shaped relationship between surprise and musical memory (**Fig. 4B**). To rule out the possibility that this result was obtained due to model complexity, we computed the Akaike Information Criterion (AIC) of the model (AIC = 1557) and compared it to a model that uses surprise as a linear predictor only (AIC = 1561). This model comparison suggests that a quadratic relationship better captures the link between surprise and memory. Overall, our findings confirm that both low- and high-surprise clips were remembered better than clips with intermediate levels of surprise.

**Figure 4:**
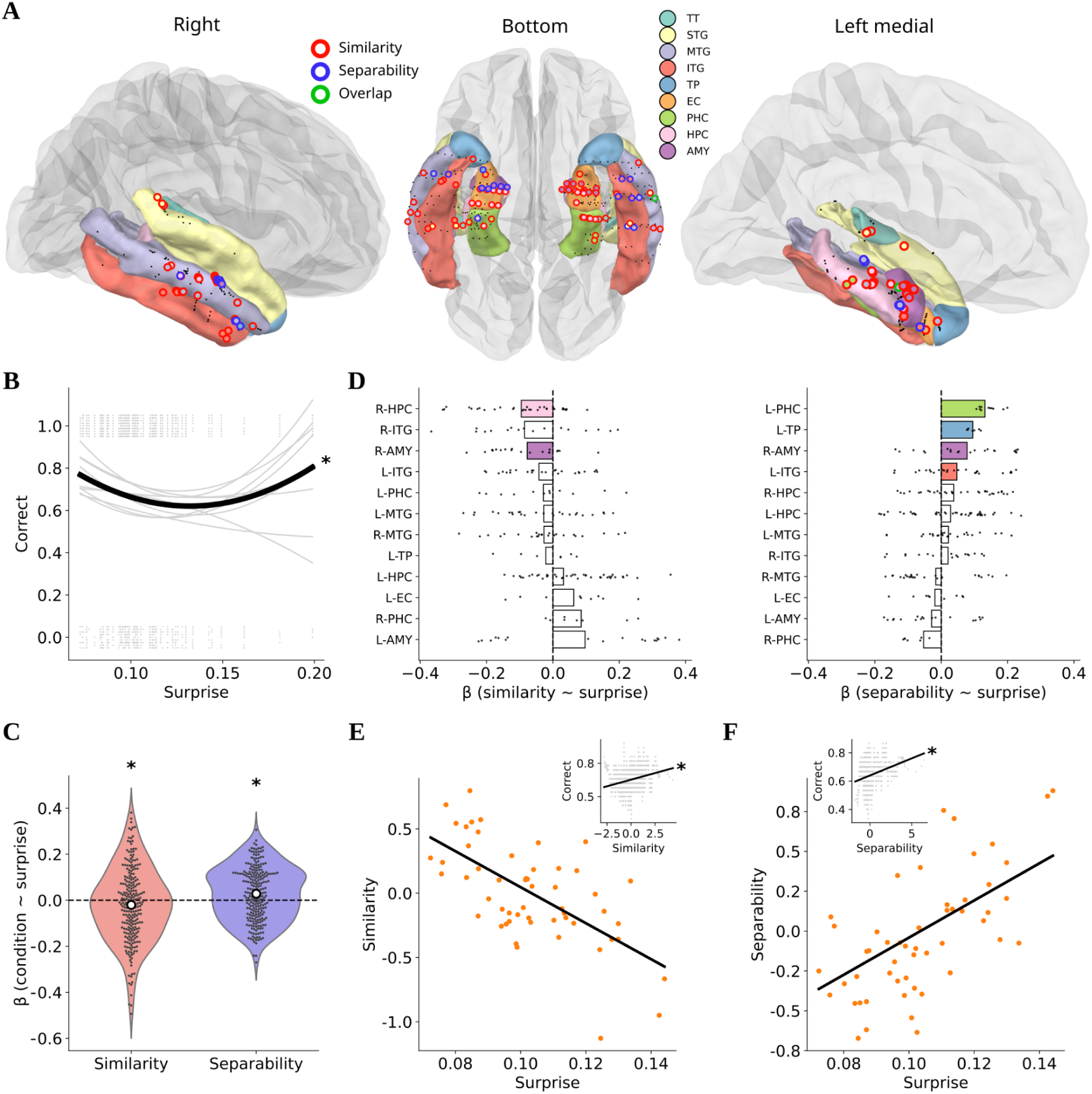
Low and high extremes of musical surprise influence neural pattern similarity and separability, respectively. **A.** Channels showing a significant effect (linear regression parameter, *p* < 0.05) of surprise on similarity (red) or on separability (blue). One channel showed both effects and is indicated in green. Inner colors indicate the anatomical location of the channels. **B.** Quadratic relationship between clip surprise and memory performance. Gray curves represent individual participants’ quadratic fits; the black curve shows the average quadratic fit, which was significant (fixed effect of squared surprise from a GLMM, * *p* < 0.05). Gray dots indicate trial-level performance (correct = 1, incorrect = 0). **C.** Violin plots of regression parameters in the ROIs (n = 249 channels) for the relationships *similarity ∼ surprise* and *separability ∼ surprise* (*t*-test, * *p* < 0.05). White dots indicate the mean. **D.** Bar plot of regression parameters by region for the two relationships (left: similarity, right: separability). Black dots represent individual channel parameters. Filled colored bars indicate regions with a significant average (negative for similarity, positive for separability, FDR-corrected *t*-tests, *p* < 0.05). **E.** Scatter plot and linear fit of similarity as a function of surprise for the red channels shown in A. Orange dots represent the average similarity across channels for a given clip (n = 54 clips). Inset: correct response as a function of surprise in the same channels and all trials. Gray dots show a rolling mean of correct over 30 data points. The black line shows the linear relationship fitted over all data points (test on Pearson’s correlation, * *p* < 0.05). **F.** Same as D, showing separability as a function of surprise for the blue channels shown in A. TT: transverse temporal, STG: superior temporal gyrus, MTG: middle temporal gyrus, ITG: inferior temporal gyrus, TP: temporal pole, EC: entorhinal cortex, PHC: parahippocampal cortex, HPC: hippocampus, AMY: amygdala, L: left hemisphere, R: right hemisphere.

### Pattern similarity and separability jointly contribute to the nonlinear effect of surprise on memory

We then tested whether neural pattern similarity and separability could account for this nonlinear effect between surprise and memory. More specifically, we hypothesized that pattern similarity would dominate at low levels of surprise, because expected stimuli enable precise neural encoding and stable responses across different presentations of the same clip. In contrast, we expected pattern separability to dominate at high surprise levels, as strong surprising events may act as critical events that distinguish a stimulus from others within the memory system. If confirmed, these two mechanisms could jointly explain the emergence of the U-shaped relationship between memory performance and surprise (see **Fig. 1E** for a schematic of this hypothesis).

To test these opposite effects of surprise on pattern similarity and separability, we fitted trial-level linear regressions predicting either similarity or separability from the average clip surprise, in each channel (p < 0.05; **Fig. 4A**). Across all channels in the ROIs, we observed a significant negative linear relationship between similarity and surprise (one-tailed one-sample *t*-test, *t*(248) = −2.01, *p* = 0.023) and a positive relationship between separability and surprise (*t*(248) = 3.85, *p* < 0.001; **Fig. 4C**), consistent with our hypothesis.

We then tested the regional specificity of such effects (one-tailed one-sample *t*-tests with FDR correction for the number of tested regions). We found a negative effect of surprise on similarity in 2/12 regions: the right hippocampus (*t*(33) = −4.04, *p* = 0.002) and the right amygdala (*t*(20) = −3.19, *p* = 0.014; **Fig. 4D**, left). We found a positive effect of surprise on separability in 4/18 regions: the left parahippocampal gyrus (*t*(11) = 9.82, *p* < 0.001), the left temporal pole (*t*(6) = 14.94, *p* < 0.001), the right amygdala (*t*(20) = 2.99, *p* = 0.015), and the left inferior temporal gyrus (*t*(25) = 2.26, *p* = 0.049; **Fig. 4D**, right). Therefore, surprise tends to decrease neural pattern similarity while increasing neural pattern separability in the temporal regions associated with memory.

To visualize these results, we plotted similarity and separability as a function of surprise for the channels showing significant effects (**Fig. 4E-F**; channels displayed in **Fig. 4A**). To ensure that selecting channels based on their surprise-related effects did not obscure their relevance for memory performance, we also plotted the probability of correct responses as a function of similarity (one-tailed Pearson’s correlation: *r* = 0.04, *p* = 0.007) or separability (*r* = 0.05, *p* = 0.008) across trials in the same channels (**Fig. 4E-F**, insets). These findings support the idea that pattern similarity and pattern separability serve as complementary neural mechanisms mediating the effect of surprise on memorability.

### Neural activity alone does not explain the nonlinear influence of surprise on memory

To verify that similarity and separability were necessary to explain the U-shaped relationship between surprise and memory (**Fig. 4B**), we asked whether neural activity alone could also explain this effect. To answer this question, we quantified the influence of clip surprise on neural activity, and specifically looked for a quadratic relationship which would suggest that neural activity mediates the effect of surprise on memory behavior. We ran a linear regression across trials in each channel of the ROIs, predicting neural activity from squared surprise (see **Methods**). We did not find a significant nonlinear effect of surprise on activity (two-tailed one-sample *t*-test across channels, *t*(248) = 1.74, *p* = 0.083, **Fig. S4**, right). In contrast, neural activity was a good predictor of memory performance across the same channels, as shown by logistic regressions (*t*(248) = 14.39, *p* < 0.001, **Fig. S4**, left), confirming the results obtained in the whole temporal lobe in the previous sections. Therefore, although trial-level activity is a relevant marker of memory, we need more specific neural mechanisms (similarity, separability) to explain the U-shaped relationship between musical surprise and memory.

### Dual model of the influence of surprise on memory

In the previous sections, we provided independent tests of our main predictions, namely that (*i*) neural pattern similarity is associated with better memory; (*ii*) neural pattern separability is associated with better memory; (*iii*) surprise decreases similarity and (*iv*) surprise increases separability. Taken together, these results support the idea that surprise influences memory through two distinct neural mechanisms, which results in a U-shaped relationship. However, our analyses were limited in two main aspects. First, we focused on linear relationships to test our hypotheses. Linear effects alone cannot explain the emergence of a U-shaped relationship between surprise and memory, as a linear combination results in a linear relationship. Second, testing the four predictions independently provided strong evidence for our hypothesis, but a mechanistic model that integrates all the intermediate effects is lacking.

In this section, we provide one plausible model of the U-shaped influence of surprise on memory performance. This model makes two specific assumptions: 1) the effect of surprise on neural pattern similarity and separability is not linear but exponential, and 2) similarity and separability are linearly combined in a logistic model to guide memory. We show that our data is compatible with this model, although other nonlinear relationships in 1) or statistical models in 2) could be tested. Once fitted, the model takes clip surprise as input and predicts memory performance (probability of correct response in that trial).

We applied our model at the channel level, in the ROIs. For each channel, we used nonlinear least squares (see **Methods**) to fit an exponential relationship between surprise and either similarity or separability (**Fig. 5A**). After this step, we selected a subset of channels based on fitting performance (see **Methods**). Then, in each channel, we used the fitted exponential curves to obtain predicted values of similarity and separability from clip surprise. Finally, we used these values together in a logistic regression that models memory performance as a function of predicted similarity and separability (**Fig. 5B**, yellow curves).

**Figure 5:**
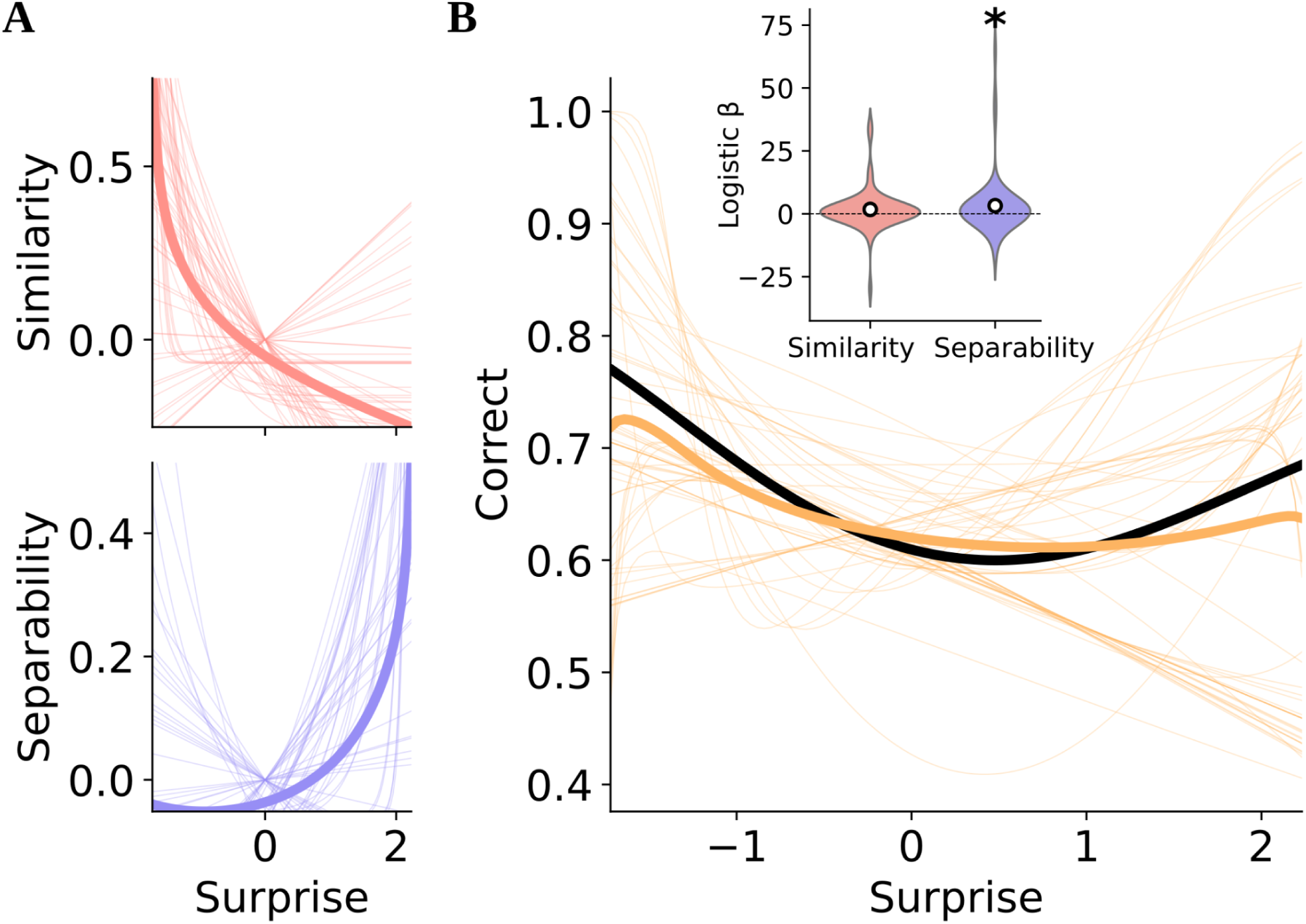
Dual model of the influence of surprise on memorability. **A.** Exponential fits between surprise and similarity or separability. Thin lines represent individual channels (n = 58) that reached the threshold in fitting performance in at least one of the two relationships, while the thick lines show the average relationships across these channels. These fits were used to obtain predicted values of similarity and separability from clip surprise. **B.** Predicted probability of correct response as a function of surprise, based on the dual model (logistic regression using predicted similarity and separability, orange) or the behavioral model (logistic regression using squared surprise, black). Thin orange lines show the dual model’s predictions in individual channels (same as A). Inset: violin plots of the logistic regression parameters of the dual model in the same channels (*t*-test, * *p* < 0.05).

To show that predicted similarity and separability are both necessary to explain how surprise influences memory, we tested the beta parameters of the logistic regression across channels (**Fig. 5B**, inset). We found a significant effect of separability on memory performance (one-sample one-tailed *t*-test: *t*(57) = 2.03, *p* = 0.023) and a trending effect of similarity (one-sample one-tailed *t*-test: *t*(57) = 1.60, *p* = 0.058). To further verify that the two mechanisms jointly influence memory, we compared our model with reduced models that use only similarity or separability. We tested the difference in model log-likelihood across channels, and found that the dual model significantly outperformed a model based only on similarity (paired two-tailed *t*-test: *t*(57) = 6.82, *p* < 0.001) or separability (paired two-tailed *t*-test: *t*(57) = 6.07, *p* < 0.001). To verify that these last results were not the direct consequence of model complexity (one additional parameter in the dual model), we applied the same procedure with models that use linear fits instead of exponential fits for the relationship between surprise and similarity or separability (see **Fig. S5**). With linear fits, the dual model did not significantly outperform the reduced models based only on similarity (paired two-tailed *t*-test: *t*(26) = −1.19, *p* = 0.878) or separability (paired two-tailed *t*-test: *t*(26) = 0.43, *p* = 0.335).

Finally, to check that our dual model provides a good mechanistic account of the emergence of the U-shaped relationship between surprise and memory performance, we compared it to a behavioral model that directly relates memory performance to surprise and is fitted at the subject level (a logistic regression with squared surprise, see **Methods**). We compared the models in terms of log-likelihood at the subject level, using the average log-likelihood across channels by subject for the dual model. We found no significant difference in model log-likelihood (L_Dual model_ - L_Behavioral model_, paired two-tailed *t*-test: *t*(8) = −0.94, *p* = 0.373; **Fig. 5B**), suggesting that our mechanistic model correctly captures what we observed in the behavior. In contrast, a dual model that used linear fits instead of exponential fits failed to capture the U-shaped relationship (L_Dual linear model_ - L_Behavioral model_, paired two-tailed *t*-test: *t*(7) = −2.52, *p* = 0.040).

Overall, these results confirm that the U-shaped relationship between surprise and memory is supported by nonlinear transformations that are applied to neural response patterns in the temporal lobe, mainly in its medial part (hippocampus, amygdala etc). Our model provides a plausible computational account of this nonlinear effect, through a combined influence of pattern similarity and separability.

## Discussion

Using a combination of computational modeling, behavioral analysis, and intracranial recordings in humans, this study reveals how statistical regularities in music excerpts nonlinearly shape memory performance via two distinct neural mechanisms. We applied a deep learning model of polyphonic musical expectations (*44*) to quantify surprise and showed that statistical unexpectedness in naturalistic classical music clips has a U-shaped influence on memory performance in an old/new task (**Fig. 4B**). Specifically, both highly expected and highly surprising clips were remembered better than those with intermediate levels of surprise.

At the neural level, we found that increased activation in higher-level regions of the temporal lobe is associated with both the detection of novel clips and the recognition of previously heard clips (**Fig. 2A-C**). Importantly, different neural populations appear specialized for novelty detection versus recognition, and these processes exhibit distinct temporal dynamics (early vs. late; **Fig. 2D-E**). We also observed a left lateralization of novelty detection (**Fig. 2A-B**), which is consistent with recent findings on novelty processing in auditory sequences (*51*). Crucially, while neural activation alone could not fully explain the complex memory effect of musical surprise, we demonstrated that two distinct neural mechanisms, pattern similarity and pattern separability, may mediate this relationship (**Fig. 3-4**). Low-surprise clips, which align closely with prior expectations, elicited more stable and reproducible neural responses across repetitions, increasing pattern similarity. In contrast, high-surprise clips enhanced pattern separability with previously heard clips, suggesting that prediction errors trigger distinct neural representations to encode new information and reduce confusion between stimuli. These two mechanisms, influenced by opposite extremes of surprise, provide a plausible mechanistic explanation for the U-shaped memory performance curve (**Fig. 4E-F**, **Fig. 5**).

Although we provided evidence for the joint involvement of neural pattern similarity and separability in memory performance, our study does not address how and where they are individually computed when we experience a stimulus. In particular, our results suggest that overlapping neural populations integrate them together to guide memory behavior, but these two distinct mechanisms are likely supported by different populations or regions (*52–54*). Likewise, we did not directly explore the computation of prediction error signals and the neural circuitry that leads to their influence on similarity and separability in the medial temporal lobe. Previous studies have provided evidence for prediction error signals during music listening (*44*, *55*, *56*), mostly in lower-level auditory regions, but have not examined how these signals shape neural representations in higher-level areas such as the medial temporal lobe. Answering these questions could inform us about the specific nonlinear transformations implemented across the processing hierarchy, to better model the relationship between predictions and memory behavior. Our dual model assumed exponential relationships between prediction errors and pattern similarity and separability, followed by a linear mapping to memory behavior (**Fig. 5**), but additional nonlinear transformations (potentially with different shapes) are likely to happen in the different steps that lead to this complex behavior.

The neural activation contrasts revealed different temporal dynamics across regions. Interestingly, although the memory task engaged both cortical and subcortical regions, early auditory areas (transverse temporal gyrus and superior temporal gyrus) did not show significant neural activation differences between correct and incorrect trials (**Fig. 2B-C**). This reinforces the idea that higher-level auditory and medial temporal regions, such as the middle and inferior temporal gyri, temporal pole, hippocampus, and amygdala, are critical for successful musical memory. Prediction errors are likely computed over the whole processing hierarchy (*57*, *58*), but only impact memory when integrated at higher cortical and subcortical levels. In line with this view, the amygdala showed a rapid, transient response to novelty (**Fig. 2E**), consistent with its well-established sensitivity to novel stimuli (*59*), while the hippocampus exhibited a slower ramping of activity, aligned with an evidence accumulation process typical of recognition (*50*). This pattern contrasts with classical episodic memory models suggesting that familiarity-based recognition is predominantly mediated by the perirhinal cortex, with the hippocampus supporting recollection (*60*). Although our task was primarily familiarity-based, hippocampal involvement was robust, indicating that recognition of musical sequences may recruit a broader temporal lobe network. Notably, our anatomical parcellation did not isolate the perirhinal cortex, a limitation that future studies should address through refined channel mapping.

Our findings align with theoretical accounts proposing that confirmed predictions facilitate memory integration, while prediction errors promote encoding of new information or the separation of memory traces (*45*). Although we did not directly assess neural integration, our results are consistent with this framework: valid predictions led to high neural similarity across presentations, while prediction errors enhanced neural separation. Future work could examine how additional stimulus repetitions modulate neural representations as a function of surprise levels.

Another important perspective of our work relates to the role of prediction errors as event boundaries, which are thought to segment continuous experience and create memory landmarks (*52*). While our analyses did not directly assess event segmentation, the observed increase in neural separability for high-surprise clips is compatible with this idea, suggesting that strong prediction errors may serve as neural markers that organize episodic memory space. It is also worth noting that our study focused on prediction errors in the context of novel naturalistic stimuli, without manipulating different types of novelty. Previous research distinguishes between contextual and stimulus novelty, each potentially influencing memory via different neural circuits and neuromodulatory pathways (*61*). Moreover, prior expectations can shape how novelty affects memory (*62*, *63*). Although our computational model of musical predictions accounted for contextual expectations, all musical stimuli were inherently novel. Future studies should explore how participants’ familiarity with stimuli interacts with prediction-based processes to refine our understanding of recognition memory.

At first glance, these findings may appear difficult to reconcile with reward-based accounts of musical memory, particularly the U-shaped relationship between surprise and memory for musical excerpts. Prior work typically reports an inverse relationship between surprise or complexity and reward-related responses (*64*), while reward itself is expected to show a monotonic relationship with memory (*65–67*). However, we did not measure musical pleasure directly, and we expect that these stimuli elicited relatively low levels of reward. The excerpts were brief, 8-second solo classical passages presented out of context and selected for consistency in note rate and note density, rather than for their capacity to evoke pleasure. PolyRNN quantifies statistical surprise, whereas musical reward additionally depends on listeners’ preferences, contextual uncertainty, and subjective valuation. Moreover, our paradigm focused on immediate discrimination among many unfamiliar excerpts, rather than on aesthetic evaluation or reward-driven consolidation. Under these conditions, low-surprise excerpts may facilitate recognition through stable, reproducible representations across presentations, whereas high-surprise excerpts may be remembered because they form more distinctive representations that reduce interference from competing traces. Excerpts with intermediate surprise may benefit less from either route. Thus, the U-shaped relationship observed here need not mirror the relationship between surprise and musical pleasure, but may instead reflect the specific computational demands of immediate recognition memory. Future work should jointly measure surprise, pleasure, and both immediate and delayed memory during music listening.

Another potential limitation concerns the scope of the stimulus set. The choice of classical polyphonic piano music may limit generalizations to a fully naturalistic environment. This choice was dictated by the use of PolyRNN, a computational model of musical expectations able to analyse naturalistic polyphonic stimuli, but in its current version only single-instrument pieces with available MIDI files (*44*). While this approach allowed us to rigorously test our hypotheses, future work could extend these methods to a broader range of musical genres and instruments to evaluate the generalizability of our findings. Advances in computational modeling may soon make it feasible to apply similar analyses to more complex and diverse musical materials.

In addition, while we examined neural dynamics using raw signal measures, future studies could incorporate finer-grained temporal and spectral analyses to clarify phase- or frequency-specific encoding processes. In light of extensive evidence showing frequency-specific oscillatory patterns and cross-frequency interactions in hippocampal memory processes (*68*, *69*), applying our approach to spectrally resolved signals may yield important insights into how information is encoded and exchanged within these structures. It would also be valuable to assess whether the neural mechanisms identified here extend to longer-term memory consolidation (e.g., overnight retention) and whether they generalize across modalities, such as speech or visual sequences. Causal manipulations (*e.g.*, targeted stimulation) could further provide direct tests of the roles of pattern similarity and separability and help clarify how medial temporal lobe regions may dynamically interact with auditory areas or broader cortical circuits to support memory encoding and retrieval.

To conclude, this study advances understanding at the intersection of memory and temporal predictive processing (*70*), demonstrating that the brain’s memory systems are shaped by the statistical structure of the environment. Our findings provide mechanistic evidence for how prediction errors and confirmatory signals jointly influence encoding through distinct neural codes, contributing to a deeper understanding of how complex auditory experiences like music are stored and recalled.

## Materials and Methods

### Participants

10 patients (3 females, age = 28.1 ± 5.4 years) with pharmacoresistant epilepsy initially took part in the study. They were implanted with depth electrodes for clinical purposes at the Hôpital de la Timone (Marseille). Neural recordings were performed between 3 to 10 days after the implantation procedure. No sedative or analgesics drugs were used, and patients received their usual antiepileptic treatment. Recordings were always acquired more than 4 hours after the last seizure. Patients were included in the study if their implantation map covered at least partially the Heschl’s gyrus (left or right). Neuropsychological assessments carried out before sEEG recordings indicated that all patients met the criteria for normal hearing. None of them had their epileptogenic zone including the auditory areas as identified by experienced epileptologists. 3 patients have had musical training (> 5 years of instrumental practice), one being a professional musician. Informed consent was obtained from all patients. The study was approved by the Assistance Publique – Hôpitaux de Marseille (health data access portal registration number PADS E2YSEB). Recordings, interpretation and analysis of SEEG were performed following the French guidelines on stereoelectroencephalography (Isnard et al., 2018) and recommendations on sEEG analysis (Mercier et al., 2022). A technical issue made the triggers unusable for half of the data of one patient. We excluded that subject from all analyses, resulting in a final sample of 9 patients for this study.

### Experimental procedure

Stimuli were chosen from the MAESTRO database (Hawthorne et al, 2019), containing midi recordings of classical piano pieces (from baroque to modern eras) performed by skilled pianists. The midi files of the entire database were analyzed in 8 second clips with step size of 1 second. Within each 8-second window, the instantaneous note rate was assessed by identifying the timing of each note onset and taking the inverse of the difference between neighboring notes, discarding intervals less than 70 ms as part of chords. After this, several features were recorded for each clip: Stability (inverse of the standard deviation of temporal intervals), Peak Frequency (most common note rate within bins of .5 Hz), Mean Frequency (mean note rate across frequencies), Onset Start (the time of the first note within the 8 second window) and Note Density (average number of concurrent notes played at each given point in time). From this information we selected the “best” clips as those with the highest Stability measurement within each Peak Frequency bin of 2.5 - 7.5 in 1 Hz steps. Only clips with an average note density between 2 and 2.5 were considered. We avoided picking clips from the same songs by ensuring the levenshtein ratio between the titles and composers of each entry were not greater than .9. Selected clips were manually inspected to remove duplicates across multiple performances which could yield the same piece due to different naming practices. The 18 most stable clips per Peak Frequency bin were chosen to be included in the experiment (108 clips in total). Remaining clips were saved as individual midi files. They were then converted to sound files using the python package Pretty Midi (Raffel and Ellis, 2014) fluidsynth function. Midi files were snapped to note onset start to ensure no extra silence at the beginning of sound files). The soundfont used to generate the audio was a recording of a Mason Hamlin piano (MasonHamlin-A-v5.2.sf2) recorded and published online for free use at Soundfonts4u (https://sites.google.com/site/soundfonts4u). For a separate planned analysis, we were interested in altering the onset shapes of the notes and therefore used the original and several altered versions of the soundfont to smooth the piano’s acoustic edge. To do so, we used the polyphone soundfont editor to manipulate the attack in 3 settings: 0.001 seconds (original setting), 0.100 seconds (smooth setting), 0.250 seconds (smoothest setting), smoothing the onset shape of the piano sound. The onset manipulation was applied to specific clips separately so that participants would not hear the same clip with different onset shapes. The application was done in a balanced way across clips so that the mean stability rank for each onset shape was controlled. The effect of onset shapes was not analyzed in the present study. The subjects completed an old/new task, where they had to indicate whether they had already heard the same excerpt during the experiment. Their response was given with a button press at the end of each trial. Overall, the experiment contained 180 trials, divided into 2 blocks of 90 stimuli. Trial order was set and frozen across all participants. It was pseudo-randomized so that repetition numbers were as balanced as possible in terms of position within the experiment. A block started with 18 trials of “new” clips, followed by a balanced presentation of “old” and “new” clips. Thus, only two-thirds of the clips were presented twice during the experiment.

### Acquisition and preprocessing

The sEEG signal was recorded using depth electrodes shafts of 0.8 mm diameter containing 10 to 15 electrode channels (Dixi Medical or Alcis, Besançon, France). The channels were 2 mm long and were spaced from each other by 1.5 mm. The locations of the electrode implantations were determined solely on clinical grounds. Patients were included in the study if their implantation map covered at least partially the Heschl’s gyrus (left or right). The cohort consists of 8 bilateral implantations and 1 unilateral implantation (right), yielding a total of 134 electrodes and 1479 channels. The data was recorded using a 256-channels Natus amplifier (Deltamed system), sampled at 512 Hz and high-pass filtered at 0.16 Hz. A monopolar reference montage setup was used for the recording. The recording reference and ground were chosen by the clinical staff as two consecutive sEEG contacts on the same shaft both located in the white matter and/or at distance from any epileptic activity. The precise localization of the channels was retrieved with a procedure similar to the one used in the iELVis toolbox (Groppe et al., 2017). First, we manually identified the location of each channel centroid on the post-implant CT scan using the Gardel software (Medina Villalon et al., 2018). Second, we performed volumetric segmentation and cortical reconstruction on the pre-implant MRI with the Freesurfer image analysis suite (documented and freely available for download online http://surfer.nmr.mgh.harvard.edu/). Third, the post-implant CT scan was coregistered to the pre-implant MRI via a rigid affine transformation and the pre-implant MRI was registered to the MNI template (MNI 152 Linear), via a linear and a non linear transformation from SPM12 methods (Penny et al., 2011), through the FieldTrip toolbox (Oostenveld et al., 2011). Based on the brain segmentation performed using SPM12 methods through the Fieldtrip toolbox, channels located outside of the brain were removed from the data (2.25%). The processing of continuous sEEG data was performed with MNE (v1.6) in a python environment. The signal was notch filtered at 50Hz and harmonics up to 250Hz to remove power line artifacts. Channels and epochs with artifacts and epileptic spikes were discarded with a semi-automatic procedure (Mercier et al., 2022). sEEG data was epoched between −2 to 8.5 seconds relative to stimulus onset, and the peak of the absolute value of the signal was taken for each epoch of each channel. For channel rejection, peak values were first averaged across epochs. Then, the mean and the standard deviations of peak values across channels were computed, and a threshold was defined as N standard deviations above the mean. The factor N was set based on visual inspection of the data (in a range from 4 to 10), and all channels with a peak value exceeding the threshold were removed from subsequent analyses. For epoch rejection, the same procedure was applied with peak values averaged across channels. This whole procedure was conducted for each patient independently, removing a total of 3 channels and 1 epoch. The following preprocessing and analyses were done in Python (v3.8.18) with MNE (v1.2.0). We baselined the epochs, removing the mean of the baseline computed in the time window (−2s, −0.1s). Finally we downsampled the signals to 400 Hz and applied a zero-phase FIR high-pass filter at 0.5Hz.

### Channel parcellation

We used an existing anatomical segmentation on the “fsaverage” subject to label the channels. We used MNE’s function mne.get_montage_volume_labels with the aparc+aseg (*46*, *47*) segmentation (distance parameter set to 2). We kept the first region that was not unknown as the label of each channel. We removed all channels in white matter, cerebrospinal fluid or unknown region from further analysis. Visualizations of 2-dimensional projections of the channels were obtained with MNE’s function mne.viz.snapshot_brain_montage. **Table S1** displays the number of channels per region.

### Model of musical expectations

PolyRNN was introduced in (*44*). It is based on a deep learning architecture composed of a Long-Short Term Memory (LSTM) layer and outputs time-resolved predictions of note probabilities (multi-label classification). It can work on naturalistic polyphonic piano sequences. The notes probability distributions of a musical excerpt can be used to extract the statistical surprise of each played note, as described in the paper. The model output was validated in (*44*), demonstrating that note surprise is encoded in auditory areas (mainly the Heschl gyrus and superior temporal gyrus). In this study, we used the average surprise of all notes as a measure of clip surprise.

### Behavioral analysis

To show the nonlinear relationship between surprise and memory performance, we used a Generalized Linear Mixed Effects Model (GLMM) to account for interindividual variability. We modeled the trial-level correctness (0 or 1), using the excerpt presentation number (1 or 2), trial number, note rate, surprise and squared surprise as fixed effects, and one random intercept per subject. We used a logit link function and a binomial distribution. Covariates were z-scored before fitting the model, and the presentation number was included as a factor.

We ran the model on R (v4.4.3) with the following formula (lme4 package):

*correct = glmer(correct ∼ presentation number + note rate + trial number + surprise + I(surprise**2) + (1|subject), data = behavior_seeg, family = binomial(link=‘logit’))*

We used the parameter of the squared surprise predictor and its associated p-value to assess the nonlinear relationship of interest. A model without squared surprise failed to explain as much variance in the data, as measured by AIC. The onset shape condition was not included in the modeling, as it did not improve the GLMM fits or yield any difference in the main results.

### Neural activity contrast analysis

Trial-level activity was computed as the root mean square (RMS) of the sEEG signal of individual channels over the clip window (8 seconds) to obtain one activity value per trial and channel. The RMS in the baseline window (−1.5s, 0s) was used to obtain a percent deviation from baseline. The RMS contrasts were based on subjects’ behavior, subtracting the average RMS of incorrect trials from that of correct ones. This was done separately in 1^st^ presentation (novelty detection) and 2^nd^ presentation (recognition).

In the RMS contrasts by region, we excluded outlier channels for visualization purposes but kept them for statistical testing. Outliers were defined as channels showing a contrast deviating by more than 3 standard deviations from the region mean.

We used a GLMM to compare the novelty detection and recognition contrasts across channels, accounting for interindividual variability with random intercepts for subjects. We included the novelty detection contrast as a fixed effect to predict the recognition contrast. Outliers (exceeding 3 standard deviations from the mean per subject) were removed from this analysis for visual purposes, but kept them to compute statistics.

The RMS contrasts in rolling time windows of one second were obtained with the same procedure as the trial contrast, computing the RMS in the windows, separately in novelty detection and recognition. The same baseline period (−1.5s, 0s) was used to obtain a percent deviation from baseline in each time window. To correct for multiple comparison and capture the temporal dependencies, the statistical test on the difference between novelty and retrieval was done across channels using a permutation temporal cluster one-sample test, implemented with MNE’s function mne.stats.permutation_cluster_1samp (10000 permutations, two-tailed). The arbitrary t-threshold to compute the clusters was chosen to set a p-value of 0.05.

To study the relationship between surprise and neural activity, we fitted a linear regression (*RMS ∼ surprise + surprise²*), using the Python package statsmodels (v0.14.0). One regression was fitted for each channel of the ROIs, including an intercept. Statistical testing was done across channels using a one-sample t-test. We used the same process to test the influence of neural activity on memory performance in the same channels, with a logistic regression (*correct ∼ RMS*).

### Similarity and separability measures

We used Pearson’s correlation as a measure of neural pattern similarity. We computed the correlation over time between the neural responses of the two presentations of a given clip, resulting in one correlation value per clip for each channel. We chose this metric to capture precise neural patterns that are stable across different presentations of a given stimulus.

We used Dynamic Time Warping (DTW) distance as a measure of neural pattern separability. DTW is a distance measure that can be used for time series evolving at different speeds (*71*). It allows the sequences to be transformed (warped) to find matching patterns that happen at slightly different times or on different time scales. We used this method to have a separability measure that accounts for imperfectly matching patterns across neural responses to similar but distinct stimuli (a property that Pearson’s correlation lacks). We used the Python library dtaidistance (v2.3.12) for the implementation. We computed the distance between the neural responses of every pair of trials, resulting in one distance matrix per channel. For each trial (row i of the matrix), we averaged its distances with all trials that had been played before (all columns j<i of the matrix) to obtain the average separability of that stimulus from the other, already heard clips. We also excluded the other presentation of the same stimulus from the average distance.

See **Fig. S6** for a visualization of the distributions of similarity and separability across trials.

As separability averages all distances with the neural responses of previous trials, the average distance of the first few trials can be biased (the sample size is small). To avoid this bias, we thus excluded the 50 first trials from the analysis. For consistency, we also removed the first 50 trials from the analysis of similarity. The impact of this choice on the relationship between surprise and memory and on the similarity and separability analyses can be visualized in **Fig. S7**.

### Similarity and separability analysis

The analysis of regression parameters was similar to the one explained in the activity contrast section. Following our hypotheses, we examined the following relationship: *correct ∼ similarity*, *correct ∼ separability*, *similarity ∼ surprise* and *separability ∼ surprise*. Relationships including similarity were done with subjects’ responses in the recognition trials only (two presentations of a given musical clip are needed). Statistical testing by region was done following the same procedure as the one on activity contrasts (one-sample t-tests with FDR correction for the number of regions), but applied to the regression parameters. Again, we excluded the outlier channels for visualization purposes only. Testing of individual channels to select the channels appearing on the brain topographies was based on individual linear (or logistic) regression p-values, with a threshold of 0.05.

The correlation between the effects of similarity and separability on correct across channels was done with a GLMM, following the procedure explained in the activity contrast section. It was also used for the relationship between the effects of surprise on similarity and separability.

Scatters of similarity or separability versus surprise were done on subsets of channels reaching threshold (p < 0.05) for the given effect. Values are z-scored by channel. The inset plots (correct versus similarity or separability) are done with the same subsets of channels, which are not selected based on correctness. The rolling means used a window of 30 data points.

To compare similarity with separability, we averaged their trial-level values across the subset of channels reaching threshold in at least one regression (*correct ∼ similarity* or *correct ∼ separability*), within each subject. We then z-scored the values across trials per subject, and concatenated all trials across subjects, obtaining 783 total trials. We used them to correlate similarity with separability across trials with a GLMM, including a random intercept per subject.

To illustrate how pattern similarity and separability jointly contribute to successful recognition, we plotted trial-level separability versus similarity and the other way around (**Fig. S3A**). The rolling means in the scatter plots used a window of 30 data points. Finally, we plotted a heatmap of the probability of correct responses in the 2D space defined by similarity and separability (**Fig. S3B**). The heatmap was constructed in four steps: (i) we binarized similarity and separability into 20 discrete intervals, (ii) we plotted the heatmap of average performance in every 2-dimensional bin, (iii) we smoothed the heatmap with a gaussian filter (standard deviation of the kernel: 1.5), (iv) we ran a cubic interpolation to fill missing bins.

### Dual model of surprise and memory

The dual model was fitted separately for each channel of the ROIs. To characterize the exponential relationship between surprise and either similarity or separability, we used nonlinear least squares fits with SciPy’s function curve_fit (v1.10.1) (*72*). Surprise, similarity and separability were z-scored before fitting. We fitted exponential curves of the form *f(x) = a * exp(b * x) + c*

We computed the coefficient of determination (R²) associated with each fit, and used this metric for channel selection. Additionally, the previous sections established a negative relationship between surprise and similarity, and a positive relationship between surprise and separability. We thus kept the channels that reached an arbitrary R² threshold of 0.05 in at least one of the two fits (*similarity ∼ surprise* or *separability ∼ surprise*) while also showing a relationship in the right direction (negative or positive, respectively), based on the sign of parameter a. See **Fig. S8** to visualize how the choice of the R² threshold can influence the results.

In the selected channels, we used the fitted curves to obtain predicted values of similarity and separability from surprise per trial. We then fitted a logistic regression to model the relationship *correct ∼ predicted_similarity + predicted_separability*, with the function LogisticRegression from the scikit-learn package (v1.3.2) (*73*).

The behavioral model was fitted by subject with the function LogisticRegression to characterize the direct quadratic relationship between surprise and memory performance: *correct ∼ surprise + surprise²*.

## Acknowledgments

This work was supported by the Fondation Pour l’Audition (FPA RD-2022-09; FPA RD-2020-10; FPA IDA012), the Agence Nationale de la Recherche under the France 2030 program (ANR-23-IAHU-0003; ANR-16-CONV-0002; ANR-24-CE17-7152-03), the Fyssen Foundation, the Fondation d’entreprise Optic 2000 - Lissac - Audio 2000, the European Union (ERC, SPEEDY, ERC-CoG-101043344) and the Excellence Initiative of Aix-Marseille University (A*MIDEX).

## Supplementary Materials

**Figure S1:**
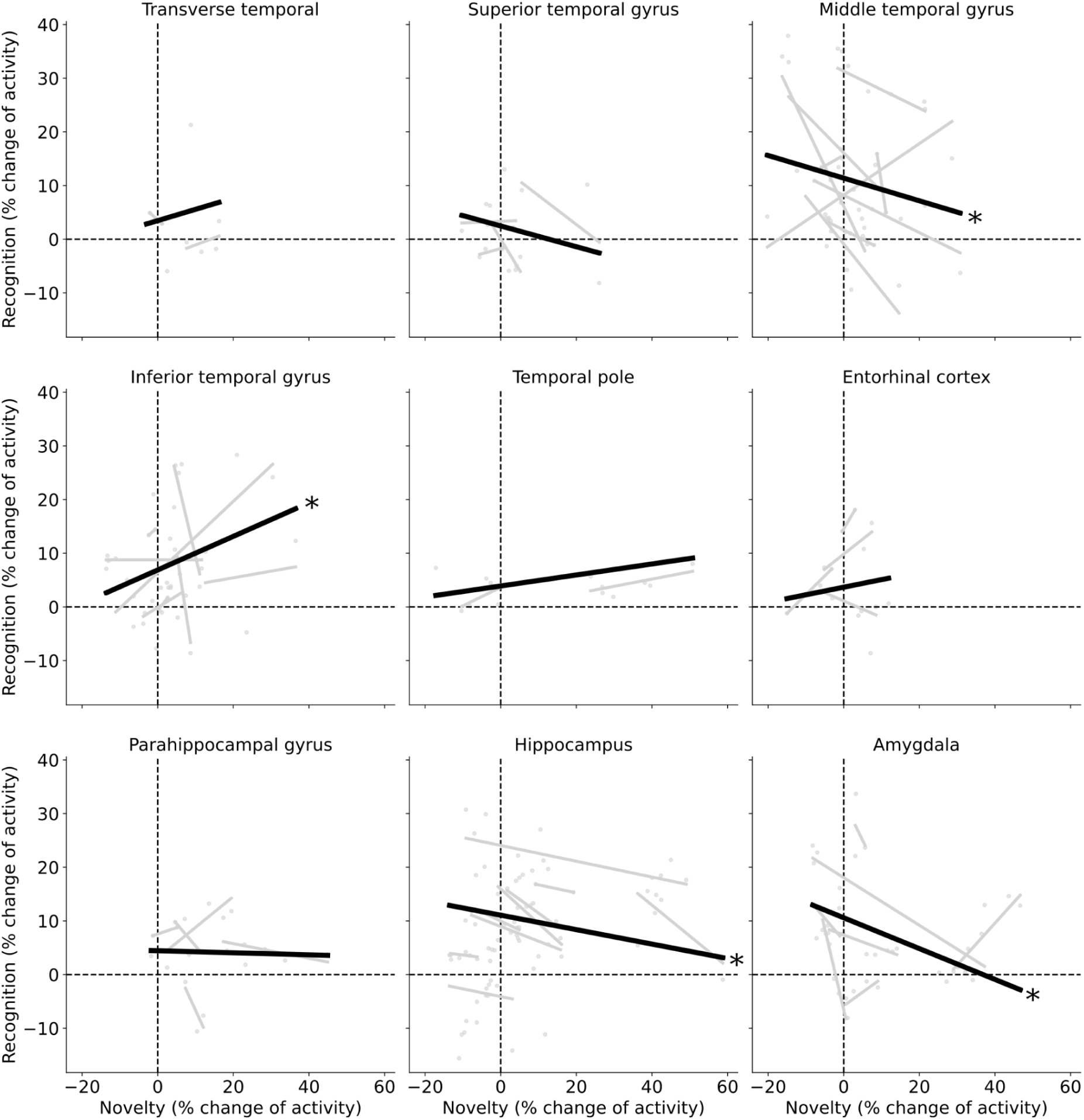
Activity contrasts in novelty and recognition trials. Recognition activity contrast plotted as a function of novelty activity contrast in each region of the temporal lobe. Gray dots/lines represent individual channels/subjects; the black line shows the average relationship (fixed effect from a GLMM, * *p* < 0.05).

**Figure S2:**
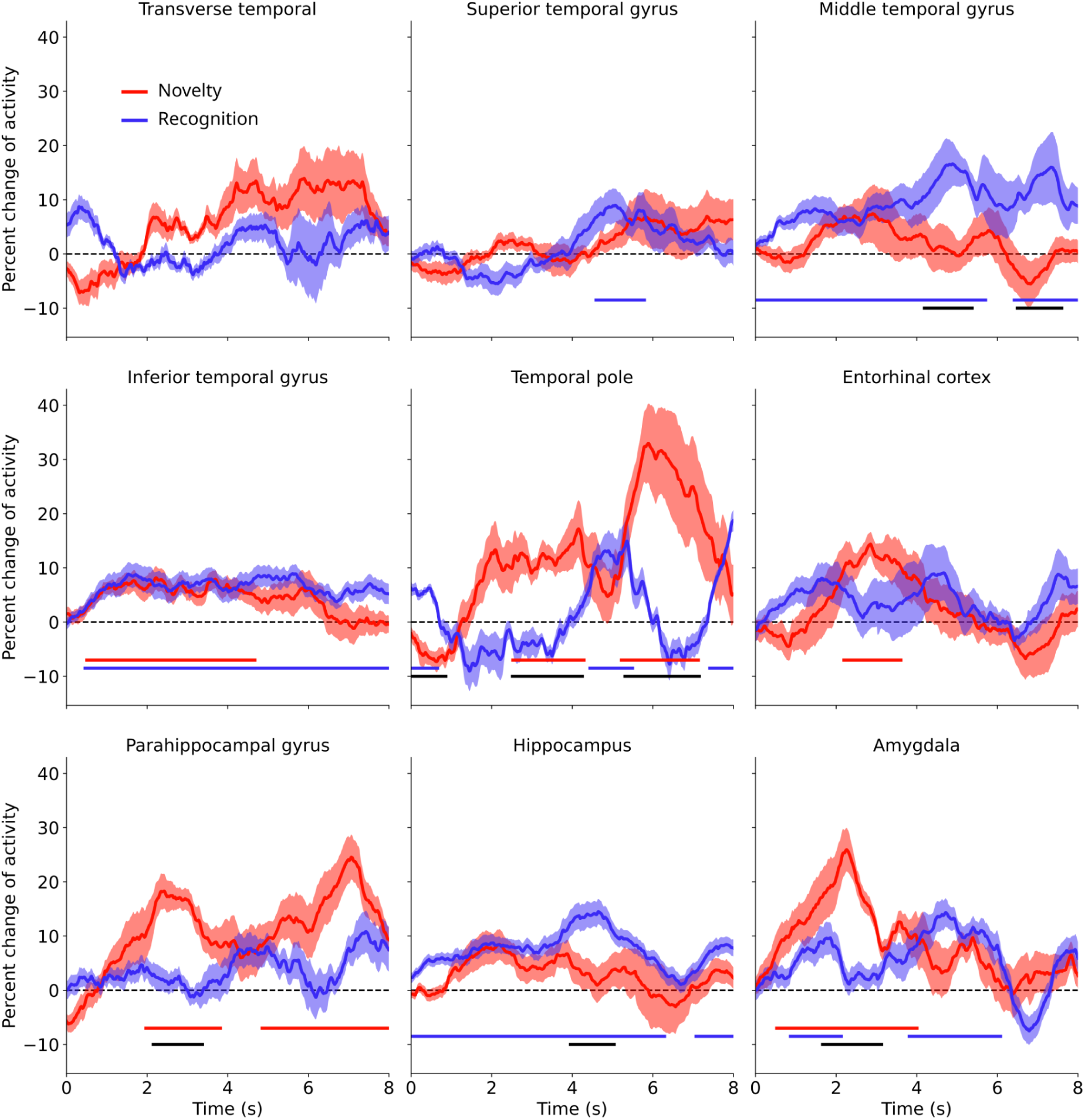
Temporal dynamics of activity contrasts. Activity contrasts over time in novelty detection and recognition, averaged across channels in each region of the temporal lobe. Shaded areas represent the SEM across channels. Lines at the bottom indicate significant temporal clusters of activity (red and blue), or differences between novelty and recognition (black), obtained from permutation-based temporal cluster *t*-tests (*p* < 0.05).

**Figure S3:**
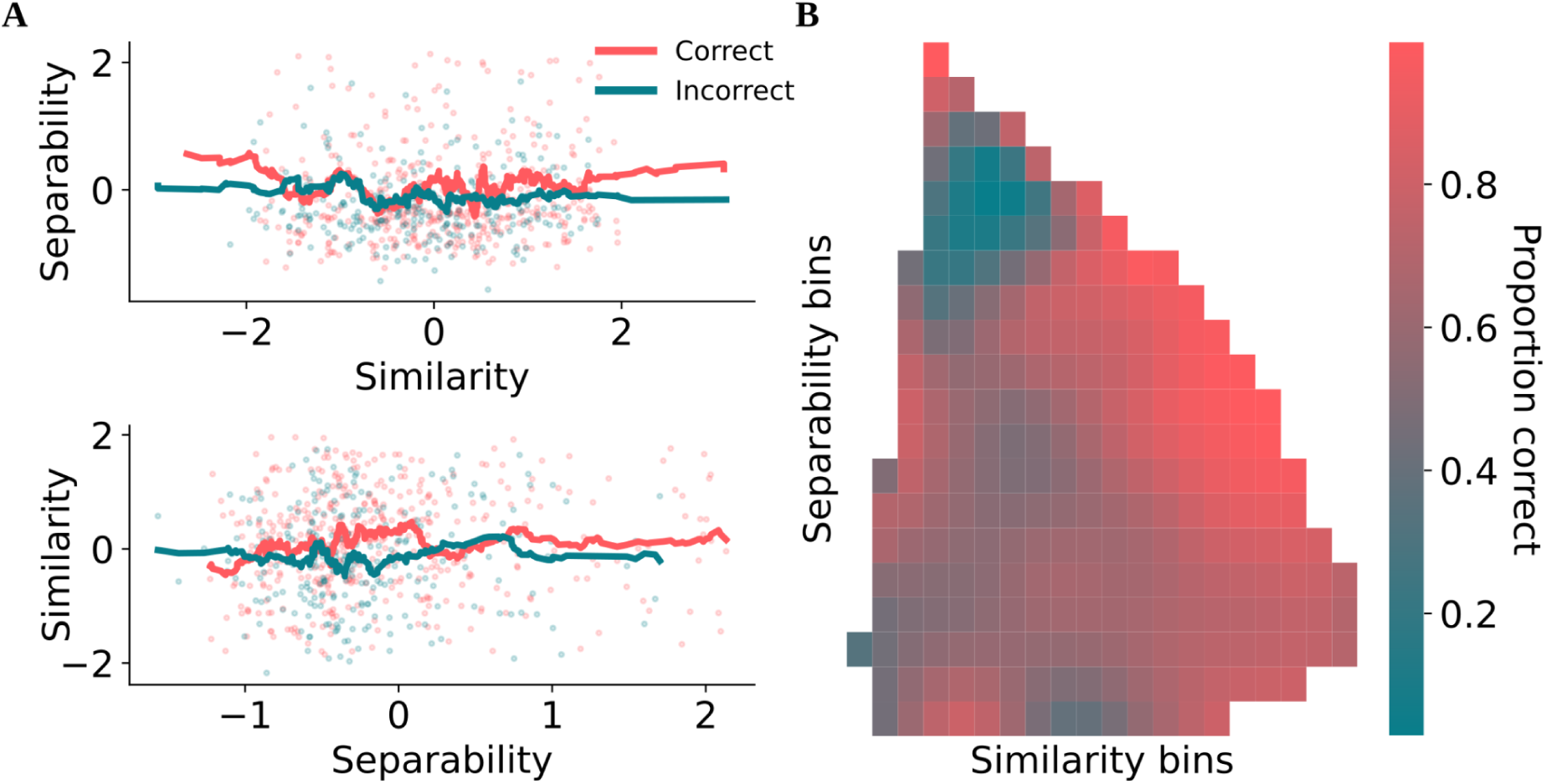
Visualization of the joint impact of similarity and separability on memory. **A.** Top: rolling mean of separability as a function of normalized (z-scored by patient) similarity, for correct (red) and incorrect (blue) trials. Bottom: rolling mean of similarity as a function of normalized separability. **B.** Heatmap of memory accuracy as a function of the normalized similarity and separability. As both similarity and separability increase, the probability of a correct response increases.

**Figure S4:**
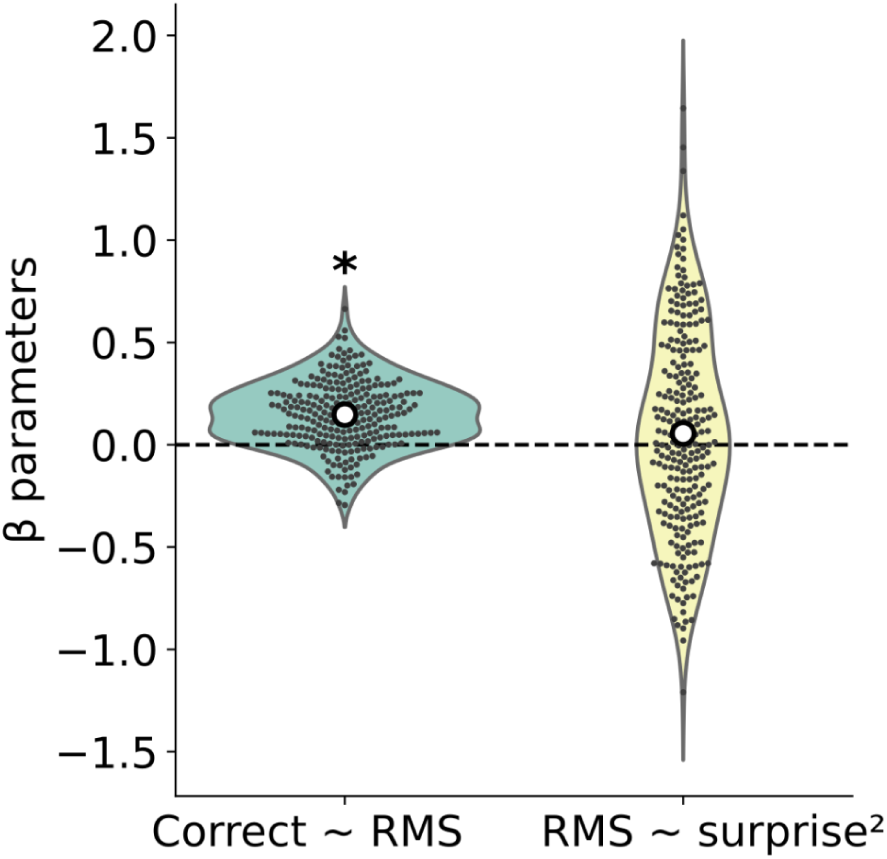
Raw neural activation cannot explain the nonlinear influence of surprise on memory. Regression parameters of the relationships *correct ∼ activity* (logistic regression, left) and *activity ∼ surprise²* (linear regression that also includes surprise as a variable, right), across the 249 channels in the ROIs. Raw neural activation, as measured by signal RMS, is associated with better memory performance but cannot explain the U-shaped influence of surprise on memory.

**Figure S5:**
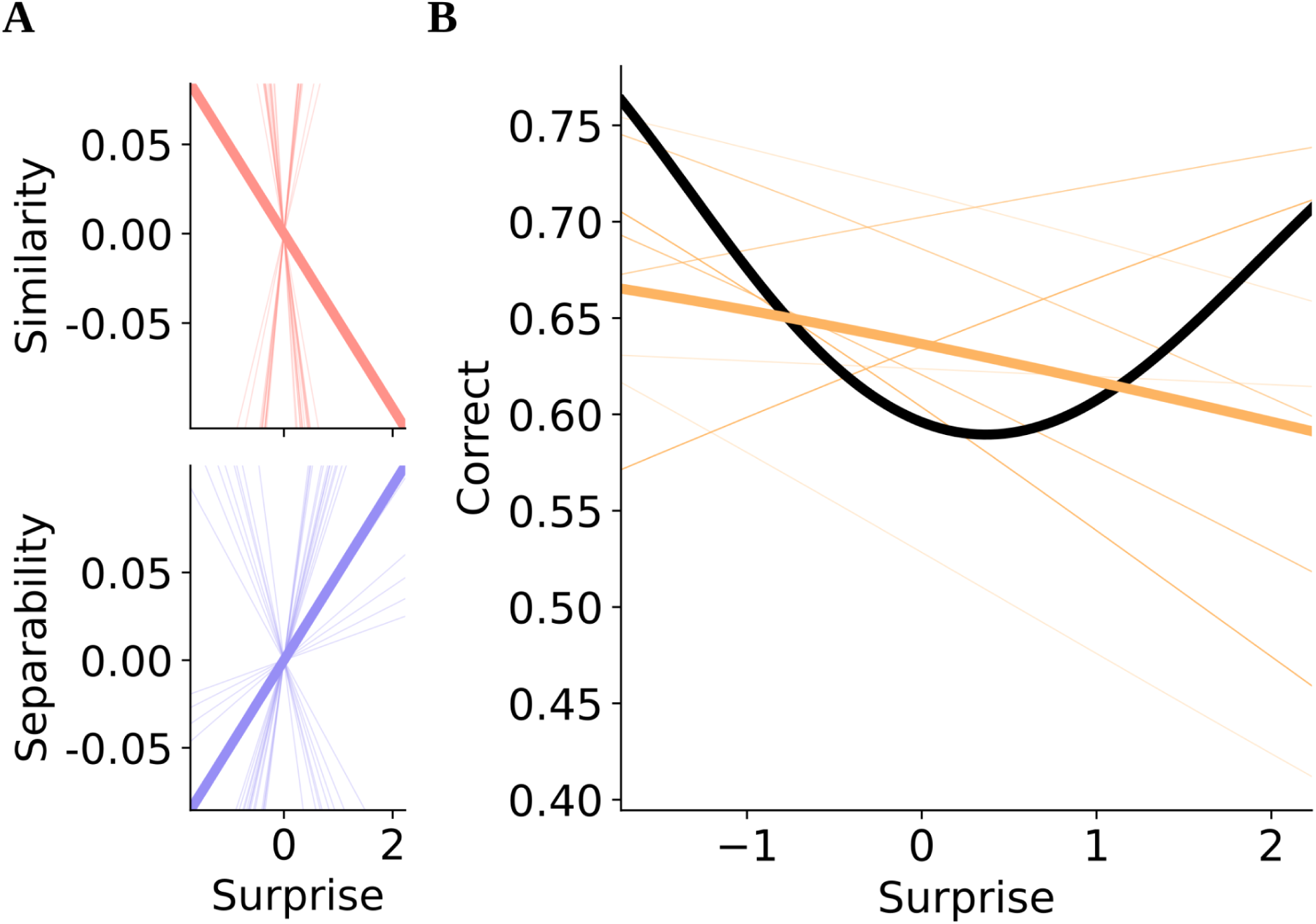
Dual model with linear instead of exponential fits. **A.** Linear fits between surprise and similarity or separability. Thin lines represent individual channels that reached the threshold in fitting performance in at least one of the two relationships, while the thick lines show the average relationships across these channels. These fits were used to obtain predicted values of similarity and separability from clip surprise. **B.** Predicted probability of correct response as a function of surprise, based on the dual model (logistic regression using predicted similarity and separability, orange) or the behavioral model (logistic regression using squared surprise, black). Thin orange lines show the dual model’s predictions in individual channels (same as A).

**Figure S6:**
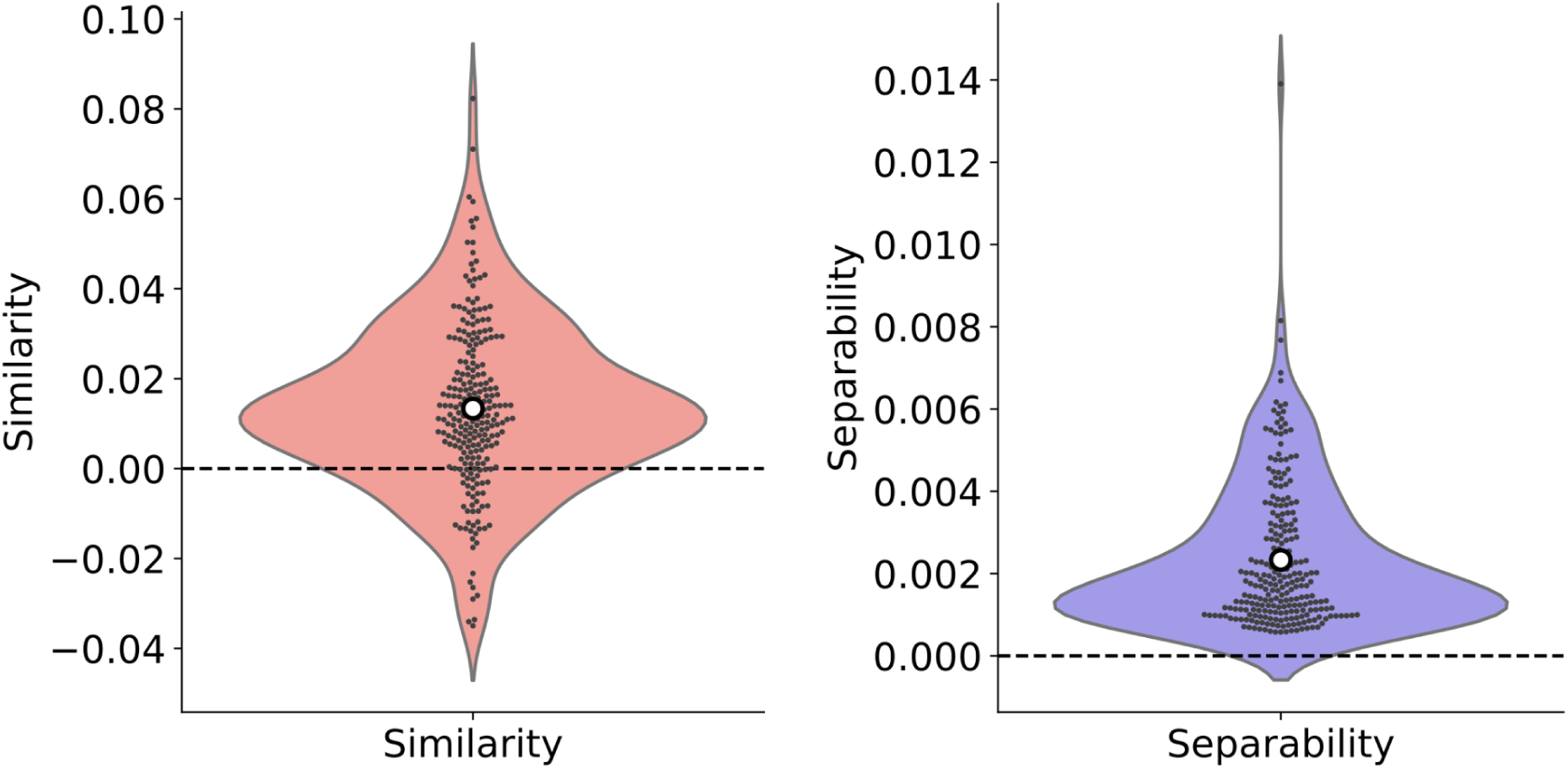
Distribution of similarity and separability. Violin plots of similarity (left) and separability (right) values from all subjects. Each dot represents a trial.

**Figure S7:**
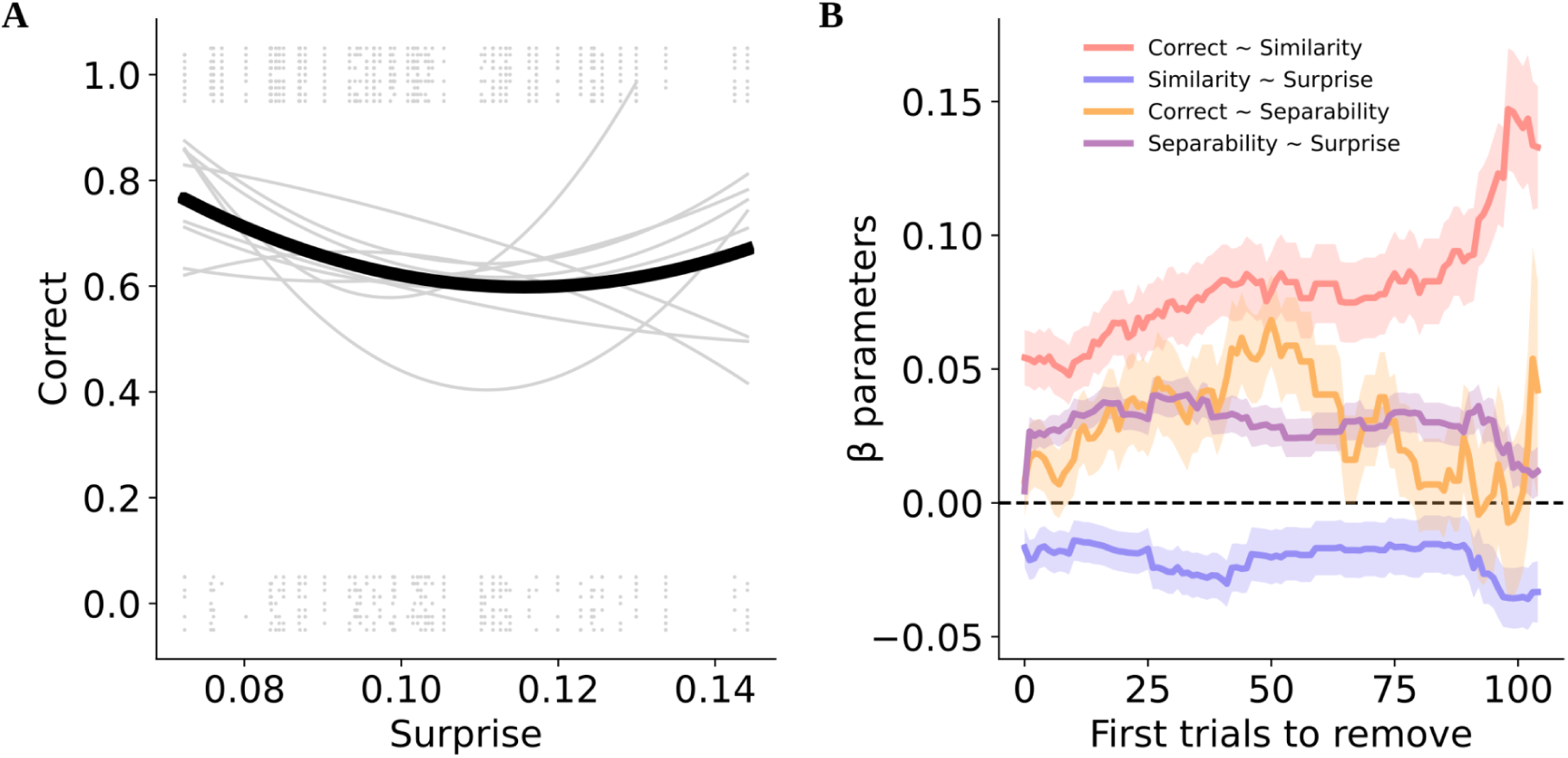
Impact of the first trials of the experiment. **A.** Behavioral results (same as Fig. 4B) without removing the 50 first trials. **B.** Evolution of the regression parameters in the four main analyses (Fig. 3D **and** Fig. 4C) as a function of the number of first trials removed from the analysis. Shaded areas indicate SEM across channels.

**Figure S8:**
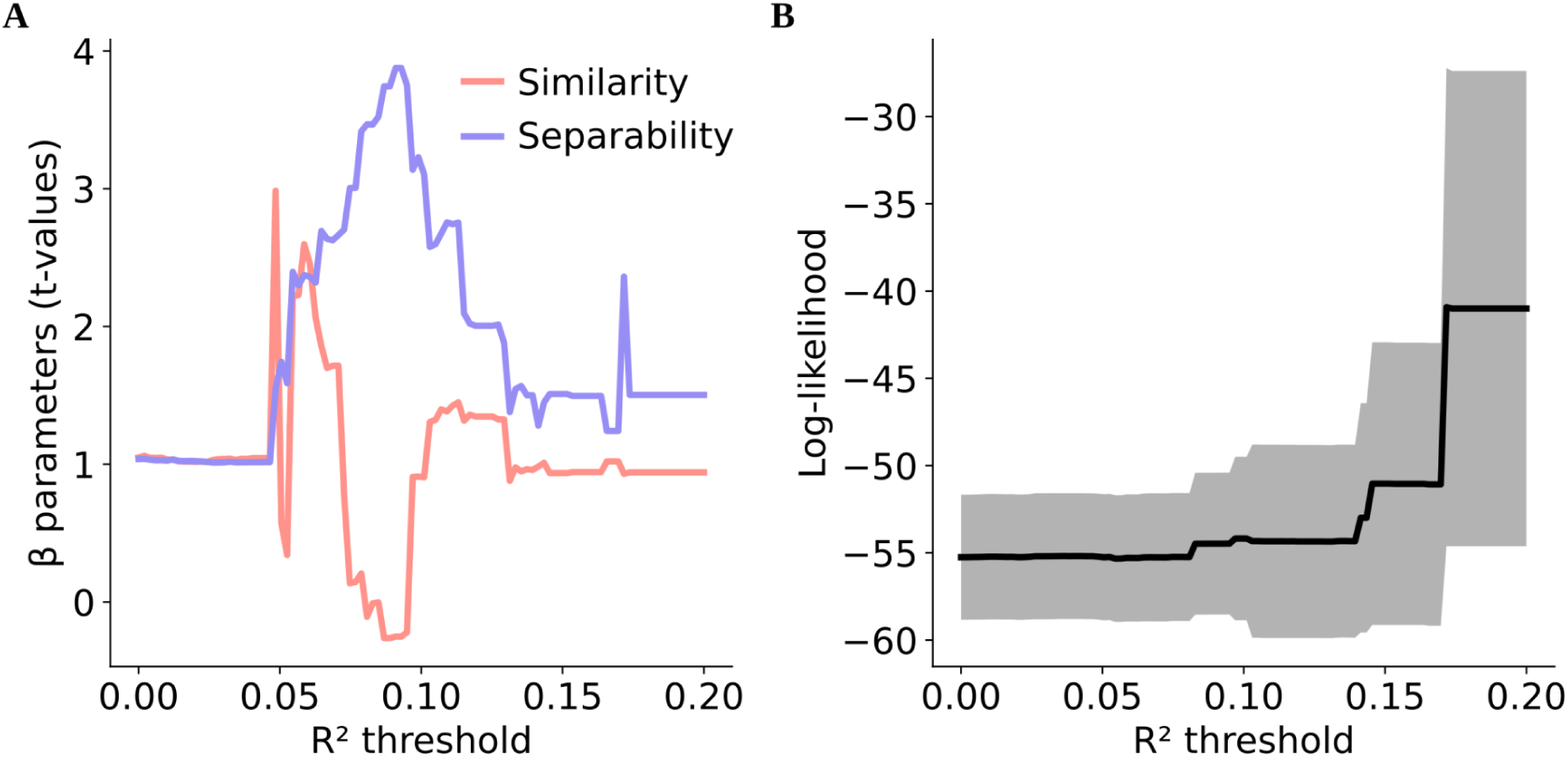
Impact of the R² threshold in the dual model. **A.** *t*-values across subjects of the logistic regression parameters, averaged across channels for each subject, of the dual model (*correct ∼ predicted similarity + predicted separability*) as a function of the R² threshold used to select channels based on exponential fit performance. **B.** Log-likelihood of the logistic regression, averaged across channels by subject. The thick line represents the mean across subjects, while the shaded area represents the SEM across subjects.

**Table S1:** Channel coverage across 9 patients. Based on the aparc+aseg segmentation (*46*, *47*).

| Region | Channel Count |
| --- | --- |
| ctx-lh-temporalpole | 7 |
| ctx-lh-middletemporal | 27 |
| Left-Amygdala | 18 |
| Left-Cerebral-White-Matter | 337 |
| ctx-lh-superiortemporal | 12 |
| Left-Hippocampus | 43 |
| ctx-lh-inferiortemporal | 26 |
| ctx-lh-insula | 9 |
| Unknown | 135 |
| Left-Thalamus-Proper | 15 |
| ctx-lh-transversetemporal | 6 |
| ctx-lh-parahippocampal | 12 |
| ctx-lh-fusiform | 5 |
| ctx-lh-lateraloccipital | 1 |
| ctx-lh-isthmuscingulate | 6 |
| ctx-lh-supramarginal | 9 |
| ctx-lh-parsopercularis | 17 |
| ctx-lh-superiorfrontal | 5 |
| Right-Cerebral-White-Matter | 443 |
| ctx-rh-middletemporal | 26 |
| CSF | 3 |
| ctx-lh-lateralorbitofrontal | 7 |
| ctx-lh-caudalmiddlefrontal | 6 |
| ctx-lh-inferiorparietal | 2 |
| ctx-lh-bankssts | 2 |
| Right-Hippocampus | 34 |
| Right-Inf-Lat-Vent | 3 |
| ctx-lh-entorhinal | 9 |
| ctx-lh-postcentral | 1 |
| ctx-rh-parahippocampal | 8 |
| ctx-rh-inferiortemporal | 18 |
| Right-Amygdala | 21 |
| ctx-rh-lateraloccipital | 1 |
| ctx-rh-inferiorparietal | 17 |
| ctx-rh-cuneus | 2 |
| ctx-rh-precuneus | 7 |
| ctx-rh-superiorparietal | 10 |
| Right-Thalamus-Proper | 13 |
| ctx-rh-transversetemporal | 3 |
| ctx-rh-superiortemporal | 9 |
| ctx-rh-supramarginal | 9 |
| ctx-rh-paracentral | 1 |
| ctx-rh-insula | 22 |
| ctx-rh-parsopercularis | 6 |
| ctx-rh-postcentral | 11 |
| ctx-rh-superiorfrontal | 3 |
| ctx-rh-caudalmiddlefrontal | 8 |
| ctx-rh-lateralorbitofrontal | 8 |
| ctx-rh-rostralmiddlefrontal | 3 |
| ctx-lh-precuneus | 5 |
| ctx-lh-superiorparietal | 1 |
| ctx-rh-fusiform | 7 |
| ctx-rh-lingual | 5 |
| ctx-rh-parstriangularis | 4 |
| ctx-rh-posteriorcingulate | 1 |
| ctx-lh-rostralmiddlefrontal | 4 |
| ctx-lh-parstriangularis | 2 |
| ctx-lh-caudalanteriorcingulate | 4 |
| Left-Caudate | 4 |
| ctx-rh-caudalanteriorcingulate | 2 |
| Brain-Stem | 1 |
| Left-VentralDC | 2 |
| ctx-rh-entorhinal | 5 |
| Right-VentralDC | 1 |
| ctx-rh-precentral | 6 |
| ctx-rh-temporalpole | 7 |
| ctx-rh-rostralanteriorcingulate | 4 |
| Right-Caudate | 3 |
| ctx-lh-parsorbitalis | 2 |
| ctx-lh-rostralanteriorcingulate | 1 |
| ctx-rh-isthmuscingulate | 1 |
| Left-Inf-Lat-Vent | 1 |

